# Seipin and Nem1 establish discrete ER subdomains to initiate yeast lipid droplet biogenesis

**DOI:** 10.1101/2020.04.20.051458

**Authors:** Vineet Choudhary, Ola El Atab, Giulia Mizzon, William A. Prinz, Roger Schneiter

## Abstract

Lipid droplets (LDs) are fat storage organelles that originate from the endoplasmic reticulum (ER). Relatively little is known about how sites of LD formation are selected, and which proteins/lipids are necessary for the process. Here, we show that LDs induced by the yeast triacylglycerol (TAG)-synthases Lro1 and Dga1 are formed at discrete ER subdomains defined by seipin (Fld1), and a regulator of diacylglycerol (DAG) production, Nem1. Fld1 and Nem1 colocalize to ER-LD contact sites. We find that Fld1 and Nem1 localize to ER subdomains independently of each other and of LDs, but both are required for the subdomains to recruit the TAG synthases and additional LD biogeneiss factors: Yft2, Pex30, Pet10, and Erg6. These subdomains become enriched in DAG. We conclude that Fld1 and Nem1 are both necessary to recruit proteins to ER subdomains where LD biogenesis occurs.

## INTRODUCTION

Lipid droplets (LDs) are evolutionary conserved fat storage organelles having multitude of functions (Olzmann and Carvalho, 2019; Walther et al., 2017). The core of LDs is comprised of neutral lipids, typically triacylglycerol (TAG) and steryl esters (SE), and is enveloped by a phospholipid monolayer. The LD periphery is decorated by structural proteins, such as perilipins, and harbors different lipid metabolic enzymes including acyltransferases and lipases. LDs are formed from the ER where neutral lipid synthesizing enzymes are located. Upon synthesis, TAG/SE accumulate between the leaflets of the ER membrane, and they coalesce into lens-like structures, that emerge toward the cytosol as they grow, while remaining connected to the ER. Accumulating evidence suggests that LD formation occurs at specific sites in ER (Adeyo et al., 2011; Chung et al., 2019; Jacquier et al., 2011; Joshi et al., 2018; Kassan et al., 2013; Salo et al., 2019; Wang et al., 2016; Wang et al., 2018). However, relatively little is known about the molecular mechanisms that control LD formation at these sites, and which proteins/lipids are necessary for the process.

Proteins implicated in LD formation include seipin (Szymanski et al., 2007), lipin (Pah1) (Adeyo et al., 2011), <u>f</u>at-storage-inducing transmembrane protein (FIT2) (Gross et al.; Kadereit et al., 2008), and Pex30 (Joshi et al., 2018; Wang et al., 2018). Seipin (BSCL2 in humans and Fld1 in yeast), is an ER membrane protein that localizes to ER-LD contact sites, and plays a crucial role in LD formation and adipogenesis (Fei et al., 2008; Salo et al., 2016; Szymanski et al., 2007; Wang et al., 2016). Mutations in human seipin result in loss of fat deposition in adipocytes, a hallmark feature of Berardinelli-Seip Congenital Lipodystrophy Type-2. Seipin facilitates maturation of nascent LDs (Wang et al., 2016), and plays a key role in TAG partitioning between small and large LDs (Salo et al., 2019). Lipin (Pah1 in yeast) encodes a phosphatidate phosphatase, which dephosphorylates phosphatidic acid (PA) to generate diacylglycerol (DAG). DAG is used by the acyltransferases, Lro1 and Dga1, to synthesize TAG, and thereby promote LD biogenesis in the ER (Adeyo et al., 2011; Barbosa et al., 2015; Karanasios et al., 2013). Deletion of Pah1 or Fld1 attenuates LD biogenesis, resulting in the accumulation of neutral lipids in the ER (Adeyo et al., 2011; Cartwright et al., 2015). FIT2, another ER membrane protein is expressed in all cell types, widely conserved and plays a crucial role in LD formation. Lack of FIT2 is lethal in higher eukaryotes (Choudhary et al., 2015; Goh et al., 2015). Budding yeast has two homologs of FIT2, Yft2 and Scs3. FIT2 proteins bind DAG and TAG *in vitro* (Gross et al., 2011), and defects in FIT2 result in aberrant emergence of LDs from the ER and the accumulation of DAG (Choudhary et al., 2018; Choudhary et al., 2015). Recently we have shown that Yft2 becomes enriched at LD biogenesis sites, colocalizes with Nem1 and Fld1, and shows co-enrichment with DAG (Choudhary et al., 2018). Two recent studies identified Pex30, a yeast protein having an N-terminal reticulon homology domain that tubulates subdomains of the ER to mark ER sites at which LDs and pre-peroxisomal vesicles originate from (Joshi et al., 2018; Wang et al., 2018). Lack of Pex30 results in a significant delay in LD formation and defects in peroxisome biogenesis (Joshi et al., 2018; Wang et al., 2018).

In this study, we show that LDs are formed from discrete ER subdomains. These sites are stable and are defined by Fld1 and Nem1. We demonstrate that the localization of Fld1 and Nem1 at these specialized ER subdomains is independent of each other, or the presence of neutral lipids, and that both are required together to create functional sites of LD biogenesis.

## RESULTS

### sLro1 localizes to sites of LD biogenesis in the ER

To visualize the earliest stages of LD biogenesis, we used a *S. cerevisiae* strain in which LD formation can be induced *de novo*. Production of neutral lipids in yeast is controlled by four enzymes: Are1, and Are2 catalyze SE production, whereas Dga1 and Lro1, synthesize TAG. Cells missing all four proteins *are1*Δ *are2*Δ *dga1*Δ *lro1*Δ (from now on called 4ΔKO) lack the capacity to produce neutral lipids, and hence have no detectable LDs (Sandager et al., 2002). All four proteins are ER-localized except for Dga1, which shows dual localization, in the ER as well as on LDs (Sorger and Daum, 2002). Expression of any one of the four enzyme is sufficient to induce LD formation (Sandager et al., 2002). Lro1, an orthologue of mammalian lecithin cholesterol acyltransferase (LCAT), has a small cytoplasmic domain, a transmembrane domain, and a large ER luminal domain containing the conserved active site residues (Fig. 1A) (Choudhary et al., 2011).

**Figure 1.**
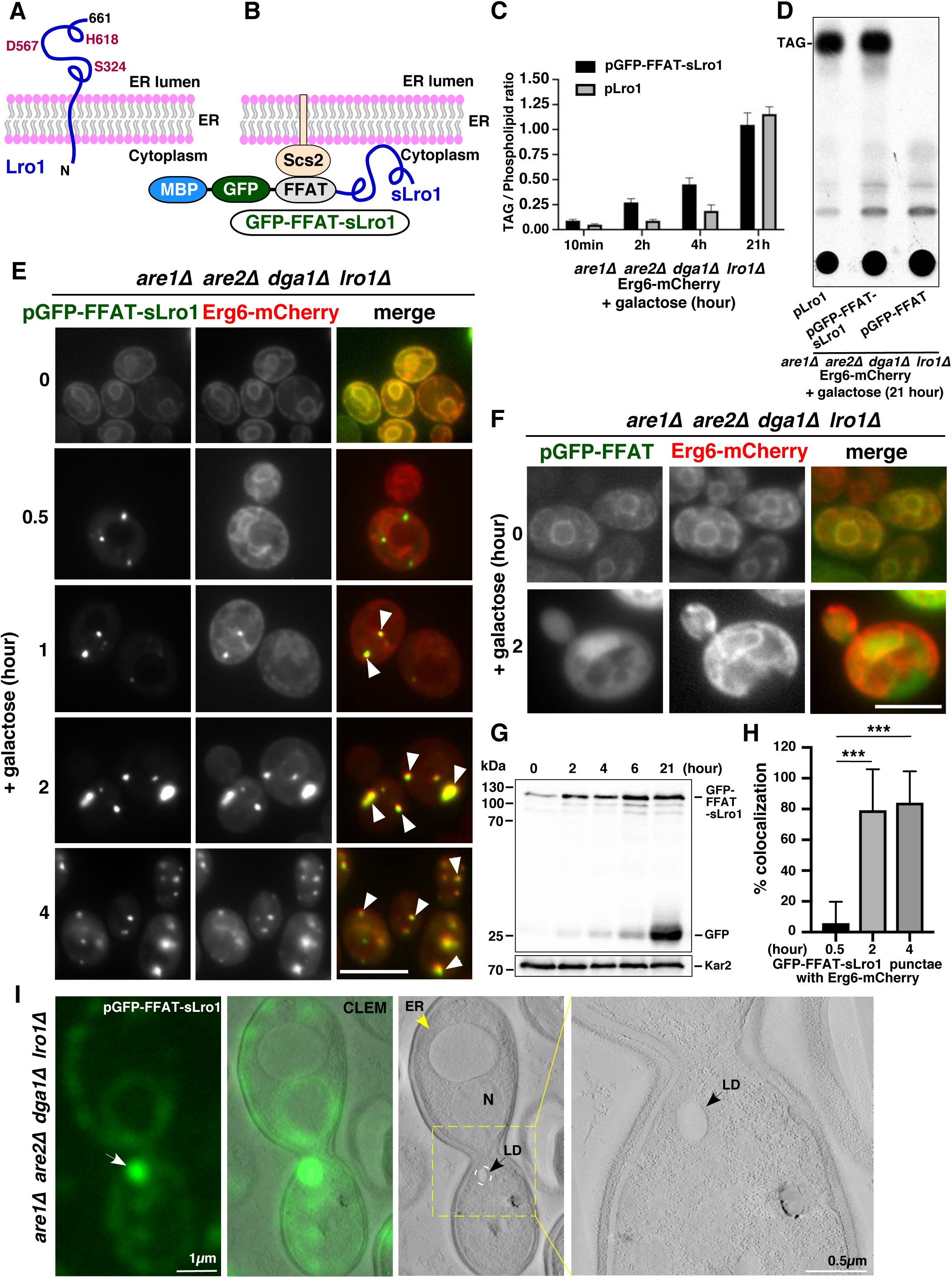
The TAG-synthase Lro1 reveals sites of nascent LD biogenesis in the ER. **A)** Topology of TAG-synthase Lro1. **B)** Design of the MBP-GFP-FFAT-sLro1 chimera, which binds the ER resident Scs2 through a FFAT motif and has its catalytic domain exposed to the cytoplasm. **C, D)** GFP-FFAT-sLro1 is enzymatically active. Cells of indicated genotypes were grown in SC raffinose, diluted in SC galactose containing [^3^H]palmitic acid, and grown for the indicated period of time. Lipids were extracted and separated by thin layer chromatography (TLC). TAG to phospholipids ratio is plotted (C), and TLC image of 21 h timepoint is shown (D). Data represent mean ± s.d. of three independent experiments. **E, F)** GFP-FFAT-sLro1 localizes to discrete ER subdomains. 4ΔKO cells expressing genomic Erg6-mCherry, and co-expressing GFP-FFAT-sLro1 (E) or GFP-FFAT (F) from a galactose inducible promoter. Arrowheads indicate colocalization between GFP-FFAT-sLro1 and Erg6-mCherry. Scale bars: 5µm. **G)** Western blot showing expression of GFP-FFAT-sLro1 or Kar2 after galactose induction for the indicated period of time. **H)** Quantification of colocalization between GFP-FFAT-sLro1 and Erg6-mCherry. Percent of GFP-FFAT-sLro1 puncta that colocalize with Erg6-mCherry after galactose induction. n > 50 cells; \*\*\**p* < 0.0005. **I)** GFP-FFAT-sLro1 colocalizes with LDs. <u>C</u>orrelative light and <u>e</u>lectron <u>m</u>icroscopy (CLEM) imaging of 4ΔKO cells expressing GFP-FFAT-sLro1. Cells were induced in galactose for 1 h. GFP-FFAT-sLro1 is enriched at a discrete site in the ER by fluorescence imaging and colocalizes with an electron translucent LD by EM. Yellow arrow demarcates ER, white arrow denotes GFP-FFAT-sLro1 puncta, black arrow denotes an LD. N, nucleus, LD, lipid droplet.

To investigate whether the soluble domain of Lro1 can catalyze TAG formation when expressed in the cytoplasm, we fused this domain of Lro1 (amino acids 98-661) to GFP, and a two phenylalanines in acidic tract (FFAT) motifs, which binds to the ER resident protein Scs2 (Loewen et al., 2003), and to the purification tag MBP (maltose-binding protein) (Fig. 1B). Scs2 is the yeast homolog of mammalian vesicle-associated membrane protein-associated proteins (VAPs), which are enriched at membrane contact sites (Wu et al., 2018). The fusion protein called GFP-FFAT-sLro1 was expressed on a plasmid under a galactose inducible promoter in 4ΔKO cells. Galactose induction of GFP-FFAT-sLro1 resulted in rapid production of TAG to levels that are similar to those observed for native Lro1, as monitored by radiolabeling cells with [^3^H]palmitic acid (Fig. 1C, D). This indicates that GFP-FFAT-sLro1 is catalytically active and can synthesize TAG in the cytoplasmic leaflet of the ER membrane.

Next, we monitored the subcellular localization of GFP-FFAT-sLro1 in 4ΔKO cells, co-expressing the LD marker Erg6-mCherry (Fig. 1E). Erg6 localizes to the ER in 4ΔKO cells, but becomes enriched on LDs as they begin to form (Jacquier et al., 2011). Upon induction in galactose containing media, GFP-FFAT-sLro1 displayed punctate localization, possibly at sites of nascent LD biogenesis, long before these sites were stained with Erg6-mCherry (Fig. 1E). By 2 h of galactose induction ∼80% of GFP-FFAT-sLro1 punctae colocalized with Erg6-mCherry (Fig. 1F). Induction of GFP-FFAT-sLro1 resulted in a time-dependent increase in protein levels as revealed by Western blot analysis (Fig. 1G). GFP fused to the FFAT motif alone, on the other hand, did not localize to sites of LD formation, or catalyze TAG formation (Fig 1F, D). Replacing the FFAT motif with the transmembrane segment of Sec63 (TMDs 1-3, aa 1-250) resulted in the same punctuate localization of the fusion Sec63(1-250)-GFP-sLro1 as observed for the FFAT containing sLro1, indicating that the puctate localization is not due to the presence of the FFAT motif (Fig. S1A, B). When induced in wild-type cells, GFP-FFAT-sLro1 localized to ER-LD contact sites and partially colocalized with Erg6-mCherry marked LDs (Fig. S1C). To determine whether sLro1 punctae correspond to sites of LD biogenesis in the ER, we performed <u>c</u>orrelative light and <u>e</u>lectron <u>m</u>icroscopy (CLEM) imaging on 4ΔKO cells. GFP-FFAT-sLro1 localized to punctate in the ER by fluorescence imaging, and these punctae correlated with electron translucent LDs in the EM sections (Fig. 1I). Taken together, these findings indicate that GFP-FFAT-sLro1 localizes to discrete sites in the ER where LDs form, suggesting that TAG synthesis is localized to specialized domains in the ER.

### TAG-synthase needs to be ER membrane anchored to produce LDs

We next asked whether the TAG-synthase, sLro1, needs to be proximal to a membrane to catalyze TAG synthesis and drive LD biogenesis. Therefore, we generated two fusion constructs of sLro1, one to express it in the cytosol, and a second to express it in the ER lumen without being anchored to the ER membrane (Fig. S1D, E). Therefore, sLro1 (98-661) was fused to GFP (GFP-sLro1), or to the signal sequence of Kar2, and a HDEL retention signal (ssKar2-GFP-sLro1-HDEL). 4ΔKO cells expressing GFP-sLro1 and ssKar2-GFP-sLro1-HDEL from a galactose inducible promoter failed to produce LDs, or catalyze any detectable TAG (Fig. S1F, G). These findings indicate that TAG synthesis by sLro1 *in vivo* requires ER membrane proximal localization of the enzyme.

### Fld1 and Nem1 localize to discrete ER subdomains even in the absence of LDs

We investigated how GFP-FFAT-sLro1 becomes enriched at discreate sites in the ER. In yeast, mature LDs mostly remain connected to the ER (Jacquier et al., 2011). Two ER proteins Nem1 and Fld1 are known to be associated with sites of LD biogenesis (Szymanski et al., 2007). Nem1 is part of a phosphatase complex (Nem1-Spo7) that activates phosphatidic acid phosphatase (Pah1), the yeast homolog of mammalian lipin, to regulate DAG formation, an important precursor for TAG synthesis (Pascual and Carman, 2013) (Fig. S2A). Nem1 shows punctate localization at ER sites that are in proximity to LDs and cells lacking Nem1 have reduced numbers of LDs, and they accumulate the neutral lipid-dye BODIPY in the ER membrane (Adeyo et al., 2011). In agreement with this, we found that the number of Erg6-mCherry marked LDs was strongly reduced in cells lacking Nem1 (*nem1*Δ), Spo7 (*spo7*Δ) or Pah1 (*pah1*Δ), and these cells displayed accumulation of BODIPY throughout the ER membrane (Fig. S2B). Like Nem1, Fld1, also localizes at ER-LD contact sites and plays an important role in maturation of LDs (Fei et al., 2008; Szymanski et al., 2007). Cells lacking Fld1 accumulate many small LDs and few supersized LDs that are up to 30-times the size of a typical wild-type LD, a phenotype that we confirm (Fig. S2C). Surprisingly, some of the BODIPY stained supersized LDs in *fld1*Δ cells failed to colocalize with Erg6-mCherry (Fig. S2C). Unlike *nem1*Δ, *spo7*Δ, *pah1*Δ or *fld1*Δ mutants, wild-type cells do not tend to accumulate neutral lipids in the ER, and LDs appear homogenous in size, as evidenced by similarly sized BODIPY stained LDs that colocalized with Erg6-mCherry (Fig. S2D). These findings suggest that the Pah1-regulating Nem1 complex and Fld1 play a role in determining the number or distribution of sites of LD biogenesis. We wondered whether the Nem1 complex plays a role in LD biogenesis that is independent of its regulation of Pah1. To test this, we expressed a constituatively active version of Pah1, called Pah1-7P, that does not require activation by Nem1-Spo7 (O’Hara et al., 2006). Cells lacking Nem1, Spo7, or Pah1 and expressing Pah1-7P showed no accumulation of BODIPY in the ER and had LDs that were similar to those observed in wild-type (Fig. S2E).

To investigate the role of Fld1 and Nem1 in determining sites of LD biogenesis, we visualized their localization in 4ΔKO cells. Surprisingly, Fld1-mCherry, and Nem1-mCherry showed punctate ER localization in 4ΔKO cells lacking any detectable BODIPY-stained LDs (Fig. S3A), indicating that the localization of Fld1 and Nem1 to discrete ER sites is independent of neutral lipid synthesis, or the presence of LDs. Next, we investigated if these Fld1, and Nem1 marked foci are stable over time. Using LiveDrop, a probe to visualize nascent LDs, seipin punctae have previously been shown to move rapidly along the ER and to stably associate LD-precursors (Wang et al., 2016). More recently, seipin foci have been shown to loose mobility upon recruitment of LD markers, LD540, LiveDrop, or the acyl-CoA synthetase ligase (ACSL3), suggesting a role of seipin in defining or stabilizing sites of LD formation (Salo et al., 2019). We visualized the mobility of Fld1-GFP, and Nem1-GFP in 4ΔKO cells by time-lapse microscopy. Some of the Fld1-GFP puncta showed rapid mobility, whereas other foci showed slow mobility (Supplementary movie 1, Fig. S3B), in agreement with previous observations (Salo et al., 2016; Salo et al., 2019; Wang et al., 2016). However, most of the Nem1-GFP punctae were stable over time (Supplementary movie 2, Fig. S3C). Overall, these data suggest that Fld1, and Nem1 localize to ER sites independent of neutral lipid synthesis, and that they likely define the location and number of potential LD biogenesis sites.

### Fld1 and Nem1 define ER subdomains that recruit the TAG-synthase

Given that the localization of Fld1 and Nem1 is independent of the presence of LDs, we wondered whether GFP-FFAT-sLro1 punctae observed upon induction (Fig. 1D), colocalize with Fld1 or Nem1. To localize sites of TAG synthesis we induced GFP-FFAT-sLro1 in 4ΔKO cells co-expressing Fld1-mCherry or Nem1-mCherry. Fld1-mCherry and Nem1-mCherry showed punctate ER localization in cells lacking LDs (Fig. 2A, B; 0 h time point). Upon induction, GFP-FFAT-sLro1 specifically localized to spots marked by Fld1-mCherry or Nem1-mCherry (Fig. 2A, B; 1 h time point). After 1 h of induction ∼80% of Fld1-mCherry (Fig. 2C, D) or Nem1-mCherry (Fig. 2E, F) punctae colocalized with GFP-FFAT-sLro1. This suggests that GFP-FFAT-sLro1 specifically localizes to sites where Fld1 or Nem1 reside, suggesting that TAG is synthesized at these discrete ER subdomains. However, in the absence of either Fld1, Nem1, or Pah1, GFP-FFAT-sLro1 failed to get recruited at these sites and instead was localized throughout the ER, suggesting that TAG is produced ectopically (Fig. 2G). This is consistent with the reported accumulation of TAG in the ER of *fld1*Δ, *nem1*Δ, and *pah1*Δ mutants (Adeyo et al., 2011; Cartwright et al., 2015). Overexpression of Fld1-mCherry in *fld1*Δ cells resulted in functional ER sites that recruited GFP-FFAT-sLro1 (Fig. 2H, Supplementary movie 3). Overall, these data indicate that ER subdomains enriched in seipin or Nem1 recruit GFP-FFAT-sLro1 to facilitate localized TAG synthesis and hence drive LD formation (Fig. 2I).

**Figure 2.**
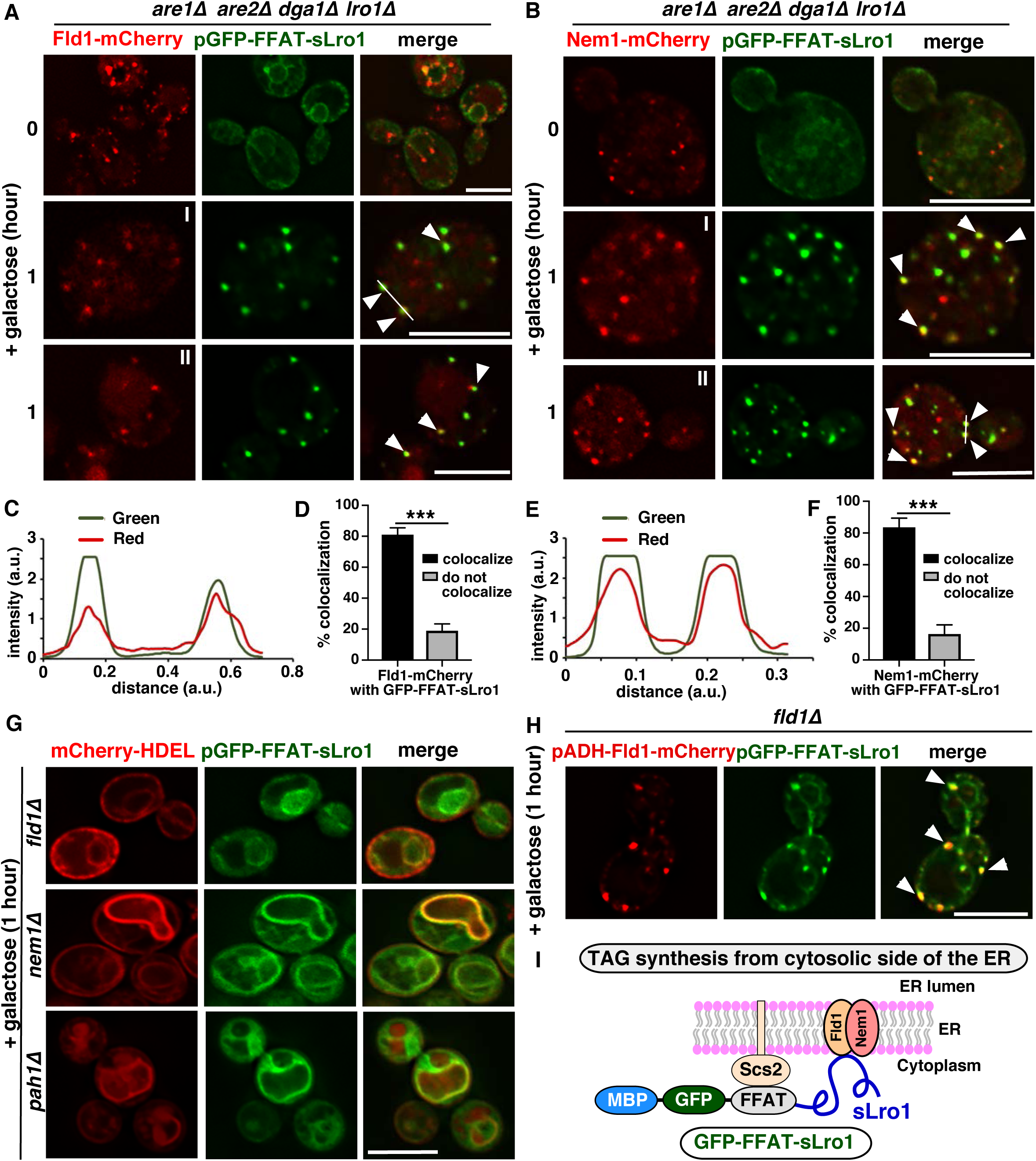
Fld1 and Nem1 define ER subdomains for the recruitment of the TAG-synthase. **A, B)** 4ΔKO cells expressing Fld1-mCherry (A) or Nem1-mCherry (B) together with GFP-FFAT-sLro1 from a galactose inducible promoter. Fld1-mCherry (A) and Nem1-mCherry (B) show punctate localization in the absence of LDs (0 h). Upon 1 h of induction, GFP-FFAT-sLro1 colocalizes with Fld1-mCherry (A) or Nem1-mCherry (B). White arrowheads indicate colocalization of sLro1 with Fld1 (A) and Nem1 (B) respectively. **C, E)** Line scan of signal intensity along the white line shown in A (I) and B (II), respectively. **D, F)** Quantification of colocalization between Fld1 (D) and Nem1 (F) punctae with sLro1. Data represent mean ± s.d., n > 50 cells; \*\*\**p* < 0.0005. **G)** Fld1, Nem1, and Pah1 are required to recruit the TAG-synthase into discrete ER foci. Cells lacking Fld1, Nem1, or Pah1 expressing GFP-FFAT-sLro1, and co-expressing mCherry-HDEL to mark the ER. **H)** Overexpression of Fld1 in *fld1*Δ cells creates ER sites that recruit GFP-FFAT-sLro1. Fluorescence microscopy of *fld1*Δ cells harboring Fld1-mCherry and co-expressing GFP-FFAT-sLro1. White arrowheads denote colocalization between Fld1 and sLro1. Scale bars: 5µm. **I)** Cartoon showing the recruitment of sLro1 at Fld1 and Nem1 sites.

### Fld1 and Nem1 recruit native Lro1

We next asked whether native Lro1 would also localize to Fld1 or Nem1 marked ER sites. Therefore, we generated GFP-tagged full length Lro1 (GFP-Lro1) and expressed it from a galactose inducible promoter. Induction of GFP-Lro1 in 4ΔKO cells co-expressing genomic Fld1-mCherry or Nem1-mCherry resulted in colocalization with Fld1 (Fig. 3A, C; 1 h time point), or Nem1 (Fig. 3B, E; 1 h time point), though GFP-Lro1 did not become as enriched in puncta as GFP-FFAT-sLro1. Upon 1 h of induction, ∼60% of Fld1-mCherry, and ∼70 % of Nem1-mCherry punctae associated with GFP-Lro1 (Fig. 3D). In cells lacking either Fld1, Nem1, or Pah1, GFP-Lro1 remained uniformly distributed throughout the ER (Fig. 3F), consistent with ectopic formation of TAG in the ER of these mutants (Adeyo et al., 2011; Cartwright et al., 2015).

**Figure 3.**
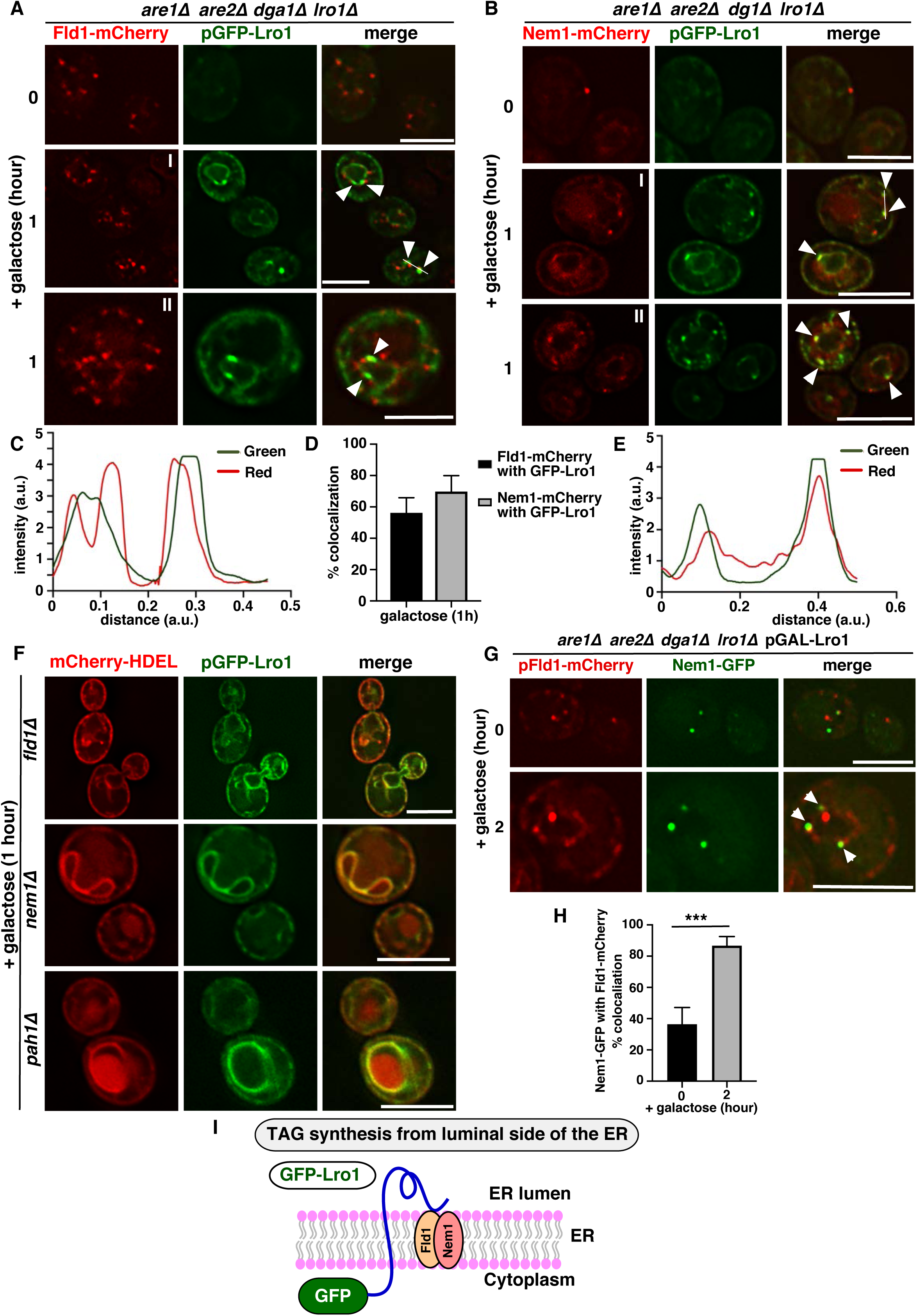
ER sites marked by Fld1 and Nem1 recruit native Lro1. **A, B)** ER subdomains marked by Fld1 and Nem1 recruit native Lro1. 4ΔKO cells expressing Fld1-mCherry (A) or Nem1-mCherry (B) and co-expressing native GFP-Lro1 from a galactose inducible promoter. Cells show punctate distribution of Fld1-mCherry, and Nem1-mCherry (0 h). Upon induction, GFP-Lro1 colocalizes with Fld1-mCherry, and Nem1-mCherry (1 h). White arrowheads indicate colocalization of Lro1 with Fld1 (A) and Nem1 (B), respectively. **C, E)** Line trace of signal intensity along the white line shown in A (I) and B (I), respectively. **D)** Quantification of colocalization between Fld1 or Nem1 foci with Lro1. Data represent mean ± s.d., n > 50 cells; \*\**p* < 0.005. **F)** Cells missing Fld1, Nem1, or Pah1 fail to recruit GFP-Lro1 into discrete foci. Cells expressing native GFP-Lro1, and co-expressing mCherry-HDEL to mark the ER, were cultured in galactose containing media for 1 h. **G, H)** Colocalization between Fld1 and Nem1 during LD biogenesis. 4ΔKO cells expressing genomic Nem1-GFP, and co-expressing Fld1-mCherry under a constitutive *ADH1* promoter, together with Lro1. Colocalization between Nem1 and Fld1 denoted by white arrowheads (G) was quantified after galactose induction of Lro1 for 2 h (H). Data represent mean ± s.d., n > 50 cells; \*\*\**p* < 0.0005. Scale bars: 5µm. **I)** Cartoon illustrating recruitment of native Lro1 at Fld1 and Nem1 sites.

To investigate how Fld1 and Nem1 organize sites of LD biogenesis, we first examined whether these two proteins colocalize. Therefore, wild-type cells expressing Nem1-GFP and co-expressing Fld1-mCherry from a plasmid were analyzed by fluorescence microscopy. Nem1-GFP showed strong colocalization with Fld1-mCherry (Fig. S3D). ∼97% Nem1-GFP punctae colocalized with Fld1-mCherry, whereas ∼33% of Fld1-mCherry was associated with Nem1-GFP (Fig. S3E). We wondered whether colocalization between Nem1 and Fld1 increases upon induction of LD biogenesis. Therefore, 4ΔKO cells expressing genomic Nem1-GFP, co-expressing Fld1-mCherry, and native Lro1 under galactose inducible promoter were analyzed. By 2 h of GAL-Lro1 induction, ∼85% of Nem1-GFP colocalized with Fld1-mCherry (Fig. 3G, H). Together, this set of results suggests that colocalization between Fld1 and Nem1 increases upon induction of LD biogenesis and that these ER subdomains recruit native Lro1 to promote localized TAG production and droplet assembly (Fig. 3I).

### ER subdomains containing Fld1 and Nem1 are enriched in DAG

Since GFP-Lro1 and GFP-FFAT-sLro1 are recruited to ER sites marked by Fld1 and Nem1, we wondered whether these sites are also enriched in DAG, because previous work suggested that sites of LD biogenesis become enriched in this lipid when LD biogenesis is induced (Joshi et al., 2018). To probe the distribution of DAG, we used a previously characterized ER-DAG sensor (Choudhary et al., 2018). This sensor is comprised of tandem C1 domains from human protein kinase D (C1a/b-PKD) fused to GFP, which in turn is fused to the transmembrane region of Ubc6, a tail-anchored ER protein (Choudhary et al., 2018). In wild-type cells the ER-DAG sensor shows ER distribution with few punctae that often colocalized with Fld1-mCherry and Nem1-mCherry (Fig. S4A). When cells were grown in the presence of oleic acid (OA) to stimulate the production of LDs, the DAG-sensor mostly localized to punctae containing Fld1 or Nem1 (Fig. 4A). Under these conditions, the ER-DAG sensor showed ∼91% colocalization with Fld1, and ∼95% with Nem1 (Fig. 4B). 4ΔKO cells also showed punctate DAG distribution similar to wild-type cells and the ER-DAG sensor colocalized with Fld1 and Nem1 sites even in the absence of LDs (Fig. S4B). However, in cells lacking Nem1, Spo7, or Pah1 the ER-DAG sensor failed to mark punctate and instead displayed uniform ER localization, probably due to altered DAG distribution throughout ther ER (Fig. S4C), suggesting that Nem1, Spo7, and Pah1 are required to produce a local pool of DAG at these ER subdomains.

**Figure 4.**
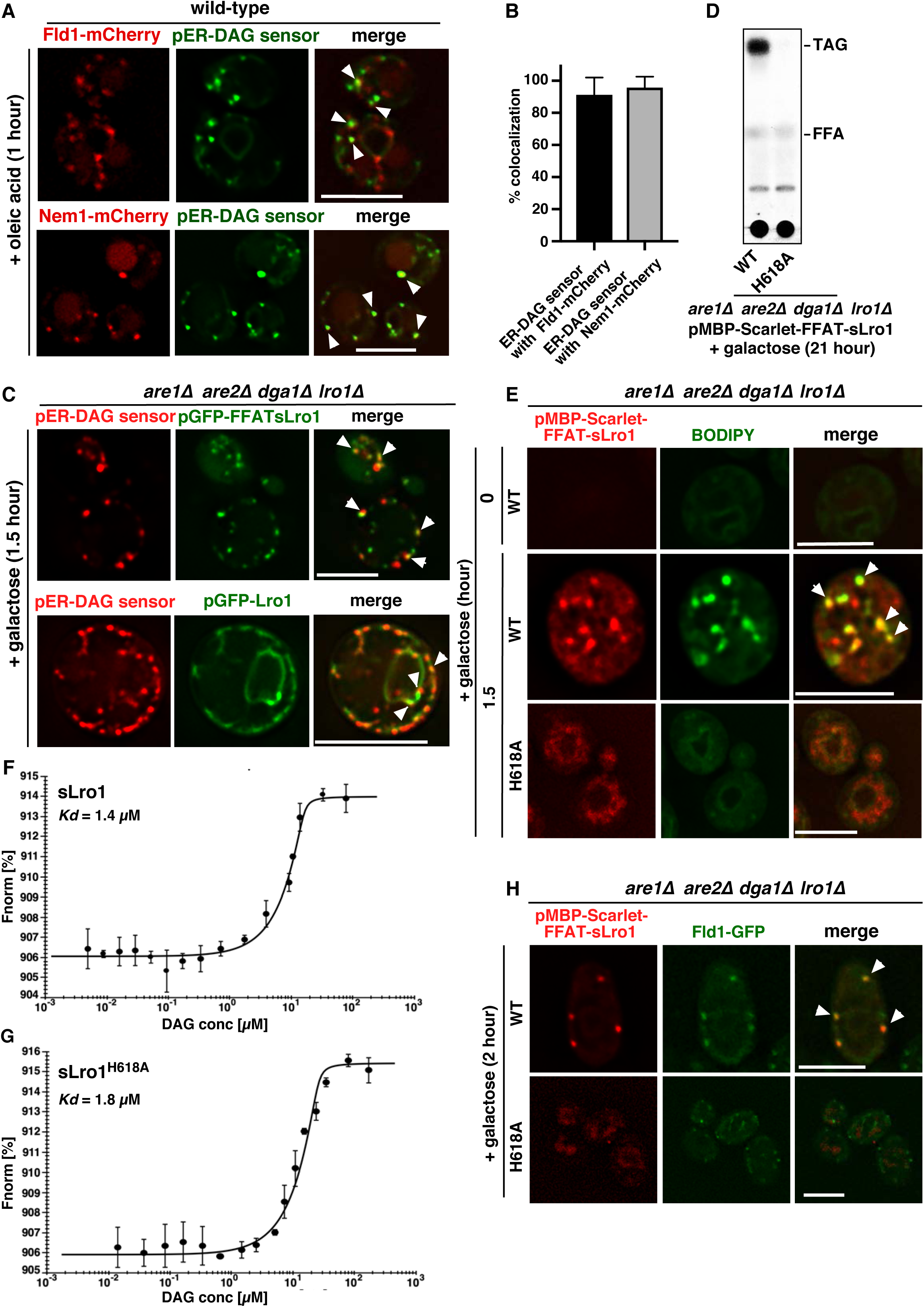
Enzymatic activity of TAG-synthase sLro1 is required for its recruitment to DAG-enriched ER subdomains. **A)** Induction of LD formation by OA results in elevated DAG levels at Fld1 and Nem1 sites. Wild-type cells expressing Fld1-mCherry, or Nem1-mCherry and co-expressing the GFP-tagged ER-DAG sensor were diluted and incubated in SC media containing 0.1% OA for 1 h. White arrowheads indicate colocalization of the ER-DAG sensor with Fld1 or Nem1 punctae. **B)** Quantification of colocalization between ER-DAG sensor punctae with Fld1 and Nem1 after 1 h OA addition. Data represent mean ± s.d., n > 50 cells; \*\*\**p* < 0.0005. **C)** Cytosolic sLro1 and native Lro1 colocalizes with the DAG sensor. 4ΔKO cells expressing mCherry-tagged ER-DAG sensor and co-expressing GFP-FFAT-sLro1/GFP-Lro1 under galactose promoter. White arrowheads denote colocalization between the ER-DAG sensor and either sLro1 or Lro1. **D)** Mutant sLro1 fails to produce TAG. 4ΔKO cells expressing WT sLro1 (Scarlet-FFAT-sLro1) or mutant sLro1 (Scarlet-FFAT-sLro1^H618A^) were grown in galactose media containing [^3^H]palmitic acid for 21 hours. Lipids were extracted and separated by TLC. TAG, triacylglycerol; FFA, free fatty acid. **E)** A catalytic inactive mutant of sLro1 fails to produce LDs. 4ΔKO cells expressing WT or mutant version of sLro1 from a galactose inducible promoter were stained with BODIPY. White arrowheads denote colocalization between BODIPY marked LDs and WT sLro1 punctae. **F, G)** WT and mutant version of sLro1 bind DAG. *In vitro* binding of the short chain DAG (C8:0) by purified WT and mutant sLro1 was measured by microscale thermophoresis. **H)** Enzymatic activity of sLro1 is required for its colocalization with Fld1. 4ΔKO cells expressing WT or mutant version of sLro1 together with Fld1-GFP were switched to galactose media for 2 h. White arrowheads denote colocalization between WT sLro1 and Fld1. Scale bars: 5µm.

To examine whether GFP-FFAT-sLro1 and native Lro1 also colocalize with the ER-DAG sensor, we examined 4ΔKO cells expressing an mCherry tagged ER-DAG sensor and co-expressing either GFP-FFAT-sLro1 or GFP-Lro1 from a galactose inducible promoter. By 1.5 h of induction, GFP-FFAT-sLro1 became highly enriched in puncta that colocalized with ER-DAG sensor (Fig. 4C). Similar results were obtained with GFP-Lro1, though it did not become as enriched in puncta as GFP-FFAT-sLro1 (Fig. 4C). Taken together, these data suggest that the TAG-synthase sLro1 and native Lro1 localize to ER sites enriched in DAG.

### Ezymatic activity of sLro1 is required for its localization to ER subdomains

Next, we wondered whether the catalytic activity of Lro1 is required for its localization to ER subdomains. Therefore, we expressed wild-type sLro1 and a catalytically dead mutant version of sLro1 (H618A) fused to MBP-Scarlet-FFAT (MBP-Scarlet-FFAT-sLro1/sLro1^H618A^) in 4ΔKO cells. When grown in glucose, cells expressing Scarlet-FFAT-sLro1 lacked any detectable LDs as revealed by BODIPY staining (Fig. S4D). Upon galactose induction, Scarlet-FFAT-sLro1 expression resulted in production of TAG as revealed by radiolabeling cells with [^3^H]palmitic acid (Fig. 4D). However, the catalytic mutant version of sLro1, Scarlet-FFAT-sLro1^H618A^ failed to form any detectable levels of TAG (Fig. 4D). This indicates that Scarlet-FFAT-sLro1 is enzymatically active and that the H618A mutant version is indeed inactive. When expressed in 4ΔKO cells, Scarlet-FFAT-sLro1 localized to punctae that stained positive for BODIPY (Fig. 4E). The catalytically inactive mutant enzyme, however, failed to generate BODIPY positive punctae (Fig. 4E). In order to gain mechanistic insight into sLro1 recruitment to ER subdomains, we purified polyhistidine-tagged sLro1 and sLro1^H618A^ from *E*.*coli* cells (Fig. S4E) and performed *in vitro* DAG binding using microscale thermophoresis. This analysis revealed that both wild-type sLro1 and catalytic dead sLro1^H618A^ bind to short chain DAG (C8:0) with comparable *Kd* values (Fig. 4F, G). Though, both sLro1and sLro1^H618A^ bind DAG *in vitro*, only the catalytically active enzyme localized to Fld1 marked ER sites (Fig. 4H). These data suggest that DAG binding alone is not sufficient for sLro1 localization to ER subdomains, but that the catalytic activity of sLro1 is required for a stable targeting to these domains.

### Fld1 and Nem1 are both required to recruit the TAG-synthase

Next, we asked whether the TAG-synthase would still be recruited to Fld1 and Nem1 containing ER subdomains when either Fld1 or Nem1 were missing. Therefore, LD formation was induced in seipin mutant cells expressing genomic Nem1-mCherry and co-expressing either native Lro1 (GFP-Lro1) or the sLro1 (GFP-FFAT-sLro1) (Fig. 5A, B). While Nem1 still localized to ER subdomains in the absence of Fld1, these Nem1 punctae failed to recruit Lro1/sLro1. These results indicate that Fld1 activity at Nem1 sites is required for recruitment of the TAG-synthase. Similarly, in *nem1*Δ cells Fld1-mCherry still localized to ER subdomains, but these Fld1 punctae failed to recruit Lro1/sLro1 (Fig. 5C, D). This set of results indicate that the activity of both proteins, Fld1 and Nem1, is required at sites of LD biogenesis for the recruitment of the TAG-synthase (Fig. 5E, F).

**Figure 5.**
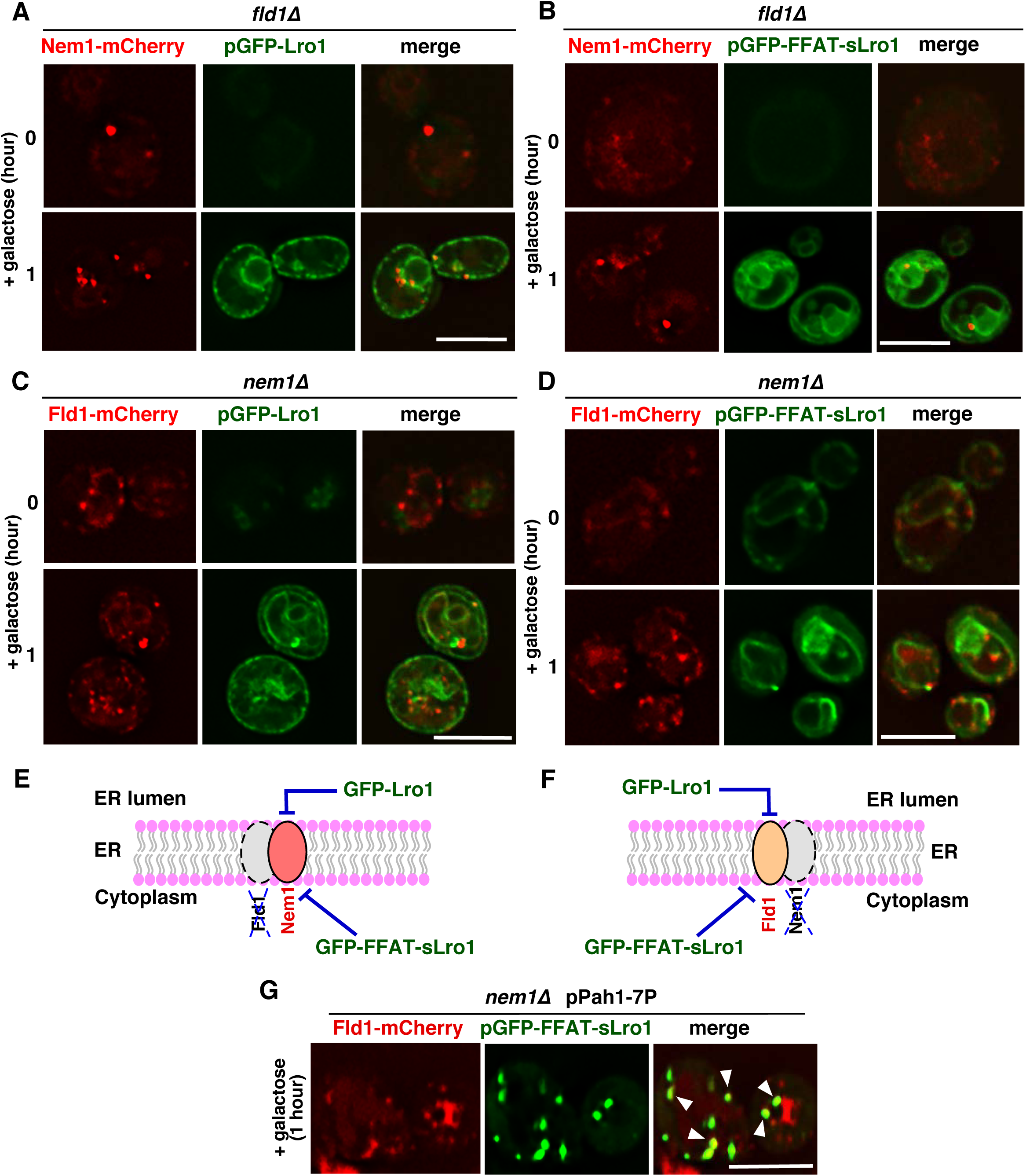
Fld1 and Nem1 are both required for recruitment of the TAG-synthase. **A, B)** Fld1 is required to recruit both native and cytosolic Lro1 to Nem1 sites. *fld1*Δ cells expressing Nem1-mCherry, and co-expressing native GFP-Lro1 (A), or cytosolic GFP-FFAT-sLro1 (B) from a galactose inducible promoter. **C, D)** Nem1 is required to recruit Lro1 to Fld1 sites. *nem1*Δ cells expressing Fld1-mCherry, and co-expressing native GFP-Lro1 (C), or cytosolic GFP-FFAT-sLro1 D) from a galactose inducible promoter. **E, F)** Fld1 and Nem1 together constitute functional ER sites for LD biogenesis. Cartoon depicting ER subdomains containing both Fld1 and Nem1 as a prerequisite for the recruitment of the native GFP-Lro1 or the cytosolic GFP-FFAT-sLro1. **G)** Expression of a hyperactive Pah1(Pah1-7P) in *nem1*Δ cells rescues the defect in recruitment of the TAG-synthase. *nem1*Δ cells expressing Fld1-mCherry, co-expressing hyperactive Pah1(Pah1-7P), and GFP-FFAT-sLro1. White arrowheads denote colocalization between GFP-FFAT-sLro1 and Fld1-mCherry. Scale bars: 5µm.

Next, we compared rates of TAG synthesis in cells lacking Fld1 or Nem1 and expressing GFP-FFAT-sLro1. Lack of Fld1 did not impair the rate of TAG synthesis when cells were labeled with [^3^H]palmitic acid (Fig. S4F). This is consistent with previous studies that found that *fld1*Δ cells had normal levels of neutral lipids (Wang et al., 2016). However, *nem1*Δ mutant cells showed a reduced rate of TAG formation and this was rescued by expression of a hyperactive allele of Pah1, Pah1-7P, which bypasses the requirement of Nem1 for activation of Pah1 (O’Hara et al., 2006) (Fig. S4F). This suggests that DAG formation in *nem1*Δ mutant cells might be limiting for recruitment of GFP-FFAT-sLro1. In support of this notion, *nem1*Δ cells expressing hyperactive Pah1 (Pah1-7P) showed normal recruitment of GFP-FFAT-sLro1 at Fld1 sites (Fig. 5G).

### ER subdomains defined by Fld1 and Nem1 recruit the LD biogenesis protein Yft2

If ER subdomains defined by Fld1 and Nem1 recruit the TAG-synthase to locally form LDs, one would expect that LD marker proteins would also co-enrich at these sites. Previously we have reported that one of the yeast FIT2 protein, Yft2, becomes enriched at sites of LD biogenesis (Choudhary et al., 2018). Yft2 together with Scs3 plays an important role in LD biogenesis, as cells missing these ER proteins have defects in LD formation (Choudhary et al., 2015). We wondered whether Yft2 would colocalize with Fld1 and Nem1. Thus, we used an inducible strain lacking three of the neutral lipids synthesizing proteins, having the fourth under a galactose inducible promoter, *GAL-LRO1 are1*Δ *are2*Δ *dga1*Δ (from now on called 3ΔKO). This strain expressed Fld1-mCherry or Nem1-mCherry and co-expressed Yft2-sf-GFP, a superfolding variant of GFP (Pedelacq et al., 2006). In cells lacking LDs, Yft2 uniformly stained the ER (Figure 6 A, B; 0 h time point). However, upon induction of TAG production, Yft2 concentrated into punctae that ∼100% colocalized with Fld1 or Nem1 (Fig. 6A-D; 1.5 h time point). Next, we determined if other ER membrane proteins would show a similar enrichment at LD biogenesis sites. We visualized the localization of the ER membrane protein, Sec63-GFP, in 3ΔKO cells expressing Fld1-mCherry. When these cells were grown in galactose media to produce LDs, Sec63-GFP did not show any co-enrichment with Fld1-mCherry marked sites of LD biogenesis (Fig. S4G), suggesting that proteins that are not involved in LD biogenesis, do not enrich at Fld1 sites.

**Figure 6.**
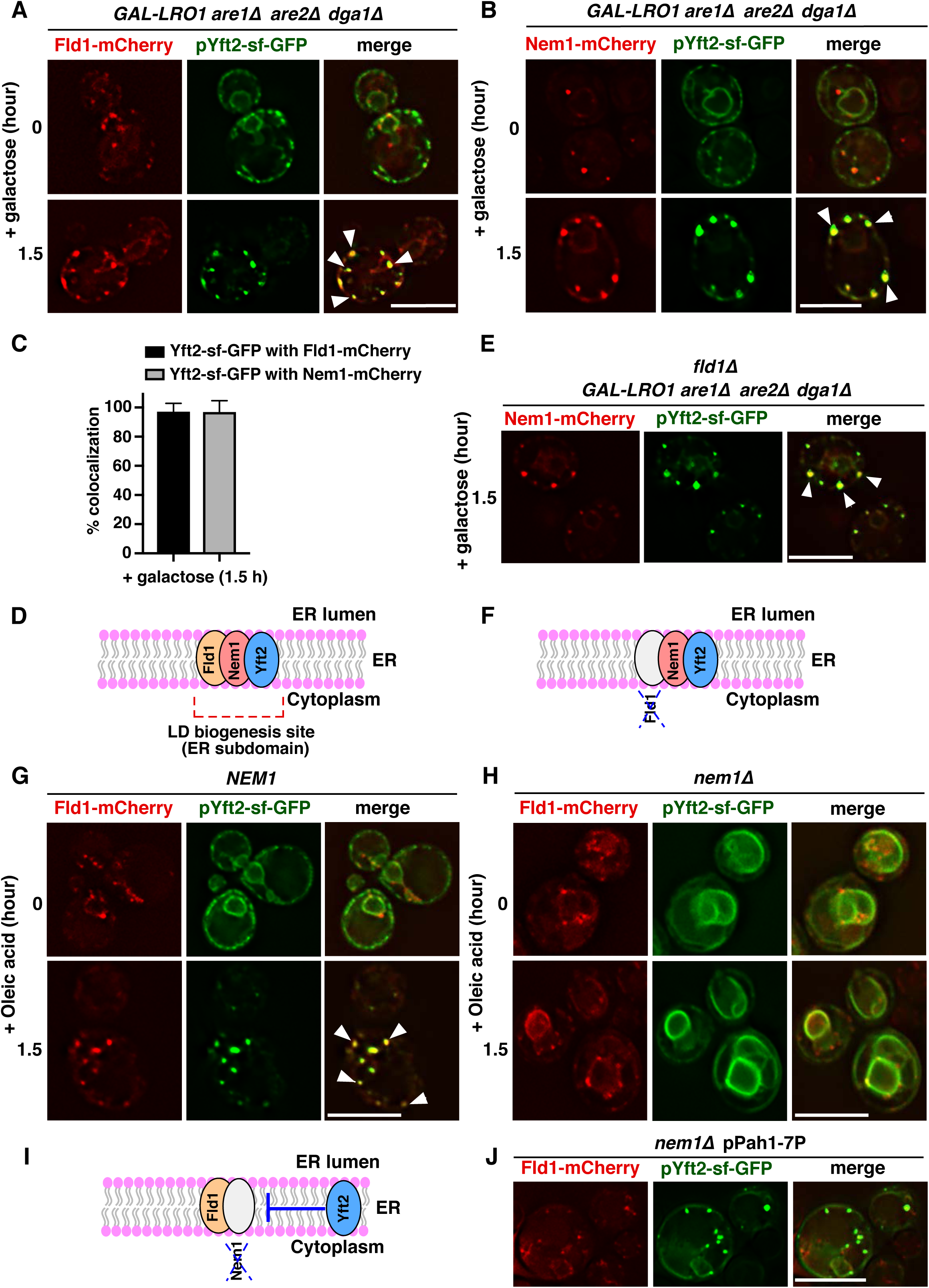
ER subdomains marked by Fld1 and Nem1 recruit Yft2. **A, B)** Yft2 concentrates at Fld1 and Nem1 containing ER sites. 3ΔKO cells expressing Fld1-mCherry (A) or Nem1-mCherry (B) and co-expressing Yft2-sf-GFP were transferred to galactose media. White arrowheads indicate colocalization of Yft2 with Fld1 (A) or Nem1 (B). **C)** Quantification of colocalization of Yft2 with Fld1 and Nem1. n > 50 cells. **D)** Cartoon depicting ER subdomain containing Fld1, Nem1, and Yft2. **E)** Localization of Yft2 at Nem1 sites is independent of Fld1. *fld1*Δ 3ΔKO cells expressing Nem1-mCherry and co-expressing Yft2-sf-GFP were grown as in (A). White arrowheads indicate colocalization of Yft2 with Nem1. **F)** Cartoon showing that the colocalization of Yft2 with Nem1 is independent of Fld1. **G, H)** Yft2 requires Nem1 at sites of LD biogenesis. *NEM1* (G) and *nem1*Δ (H) cells expressing Fld1-mCherry and co-expressing Yft2-sf-GFP were grown in SC media, resuspended into fresh media containing 0.1% OA, and visualized after 1.5 h. White arrowheads denote colocalization of Yft2 with Fld1 foci in G. Scale bars: 5µm. **I)** Cartoon depicting colocalization between Yft2 and Fld1 is dependent on Nem1. **J)** DAG levels affect the localization of Yft2 and Fld1. *nem1*Δ cells expressing Fld1-mCherry, Yft2-sf-GFP, and co-expressing hyperactive Pah1(Pah1-7P) were grown to early stationary phase in SC media. White arrowheads denote colocalization of Yft2 with Fld1 foci. Scale bars: 5µm.

### Yft2 requires Nem1 but not Fld1 at LD biogenesis sites

To determined whether lack of Fld1 from LD biogenesis sites would impair recruitment of Yft2, we generated *fld1*Δ 3ΔKO cells expressing Nem1-mCherry and Yft2-sf-GFP. Lack of Fld1 did not abrogate the recruitment of Yft2 at Nem1 sites upon induction of TAG synthesis (Fig. 6E), suggesting that the yeast FIT2 protein, Yft2 does not require Fld1 to get recruited at LD biogenesis sites (Fig 6F). Previously we have shown that Yft2, and Scs3 regulate DAG levels at sites of LD biogenesis in the ER (Choudhary et al., 2018). Since Nem1 regulates DAG production, we wondered whether Yft2 enrichment at LD biogenesis sites is dependent on Nem1. In *NEM1* wild-type cells expressing Fld1-mCherry and Yft2-sf-GFP, Yft2 showed uniform ER staining at the beginning of OA stimulated induction of LD biogenesis. However, after 1.5 h, Yft2 colocalized with Fld1 (Fig. 6G). When we performed similar OA supplementation experiments in *nem1*Δ cells, Yft2 did not enrich at discrete ER sites (Fig. 6H). Though, deletion of Nem1 did not affect the localization of Fld1, the recruitment of Yft2 to Fld1 subdomains depends on Nem1 (Fig. 6H, I). In agreement with this, expression of the hyperactive allele of Pah1, Pah1-7P, rescued the recruitment of Yft2 at Fld1 sites in *nem1*Δ cells (Fig. 6J). These data indicate that DAG levels affect the colocalization of Yft2 and Fld1, and support the idea that both Fld1 and Nem1 are needed to establish ER subdomains for LD formation.

### Lack of FIT2 does not abrogate the recruitment of the TAG-synthase

Next, we investigated whether FIT2 proteins, Yft2, and Scs3 are necessary for the localization of GFP-FFAT-sLro1. Hence, we tagged Fld1-mCherry, in *yft2*Δ *scs3*Δ double mutants expressing GFP-FFAT-sLro1. Lack of FIT2 proteins had no effect on the localization of Fld1, and these Fld1 subdomains were functional in recruiting GFP-FFAT-sLro1 (Fig 7A). Since Fld1 and Nem1 mark ER sites of LD biogenesis, and Yft2 requires Nem1 to get recruited at Fld1 sites (Fig. 6 G-J), we evaluated the distribution of DAG in *yft2*Δ *scs3*Δ double mutant cells. Therefore, we used wild-type and *yft2*Δ *scs3*Δ cells expressing Fld1-mCherry and co-expressing the ER-DAG sensor. Unlike wild-type cells where the DAG probe shows uniform ER distribution with a few spots that colocalized with Fld1, *yft2*Δ *scs3*Δ mutant cells showed large blobs containing both the ER-DAG sensor and Fld1 (Fig. 7B). These results indicate that FIT2 activity is required downstream of Fld1 and Nem1 for proper DAG distribution at sites of LD biogenesis (Fig. 7B, C).

**Figure 7.**
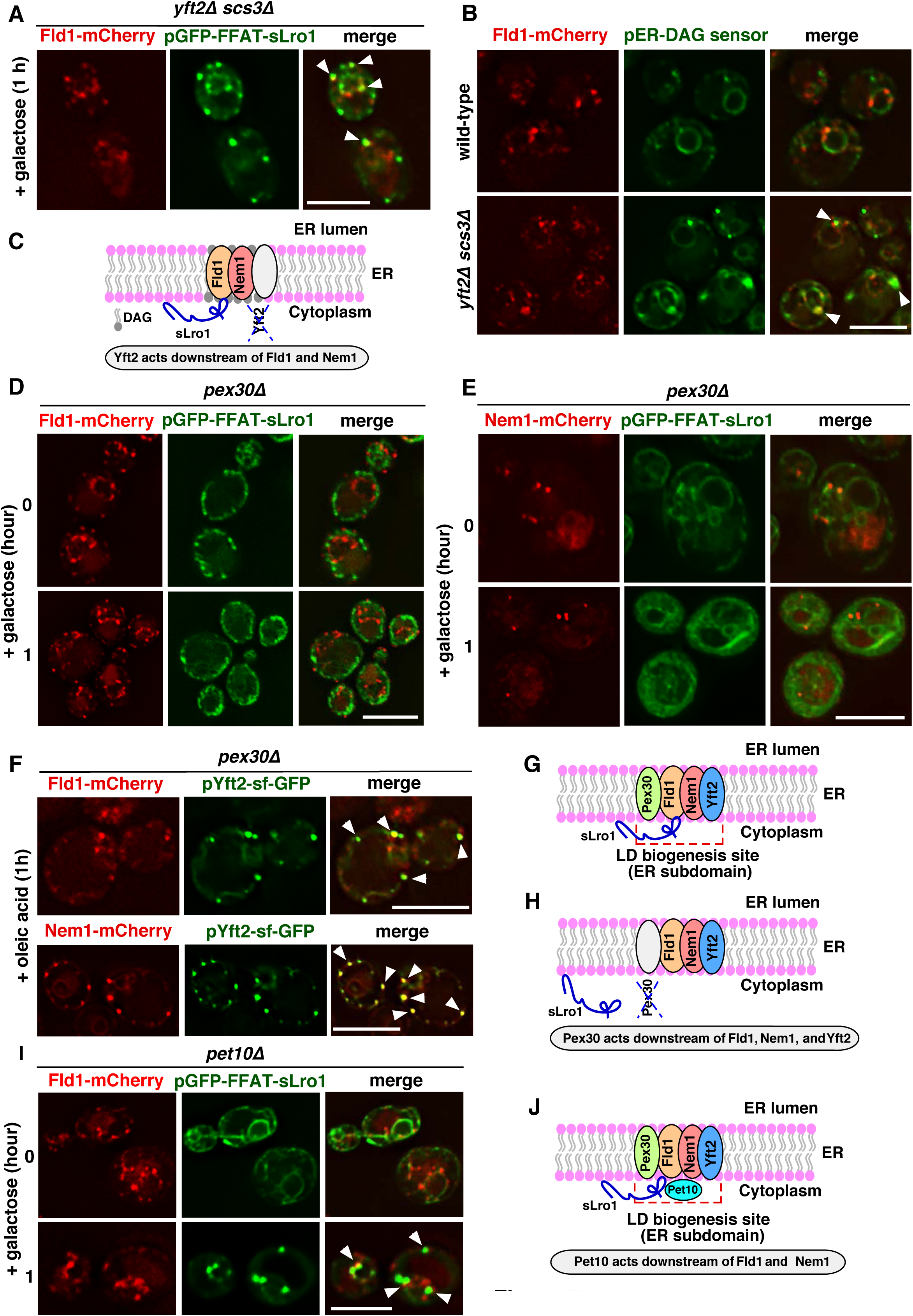
Epistatic relation of proteins at LD biogenesis sites. **A)** Lack of FIT proteins does not abrogate recruitment of GFP-FFAT-sLro1 at Fld1 sites. *yft2*Δ *scs3*Δ double mutant cells expressing Fld1-mCherry and co-expressing GFP-FFAT-sLro1 were switched to galactose media for 1 h. White arrowheads indicate colocalization of Fld1 with sLro1. **B)** Cells lacking FIT proteins have elevated DAG levels at Fld1 sites. Wild-type and *yft2*Δ *scs3*Δ double mutant cells expressing Fld1-mCherry and co-expressing the GFP-tagged ER-DAG sensor were grown to early stationary phase. Arrowheads indicate colocalization of Fld1 foci with the DAG-sensor punctae. **C)** Cartoon showing accumulation of DAG at Fld1 and Nem1 marked ER subdomains when these sites are missing Yft2. **D, E)** Pex30 acts downstream of Fld1, and Nem1 in recruiting GFP-FFAT-sLro1 to sites of LD biogenesis. *pex30*Δ mutant cells expressing Fld1-mCherry (A) or Nem1-mCherry (B) and co-expressing GFP-FFAT-sLro1 were switched to galactose containing media for the indicated time. **F)** Yft2 functions independently of Pex30 at sites of LD biogenesis. *pex30*Δ mutant cells expressing Fld1-mCherry or Nem1-mCherry and co-expressing Yft2-sf-GFP were diluted into fresh SC media containing 0.1% OA, and imaged after 1 h. White arrowheads indicate colocalization of Yft2 with Fld1 or Nem1 punctae. **G, H)** Cartoon illustrating the localization of Fld1, Nem1, Yft2, and Pex30 at ER subdomains to constitute functional LD biogenesis sites (G). Absence of Pex30 from these sites does not affect the localization of Fld1, Nem1, and Yft2, but these sites fail to recruit the sLro1 (H). **I)** Lack of Pet10 does not impair the recruitment of GFP-FFAT-sLro1 at Fld1 sites. *pet10*Δ mutant cells expressing Fld1-mCherry and co-expressing GFP-FFAT-sLro1 were grown as in D. Arrowheads denote colocalization of sLro1 with Fld1 foci. Scale bars: 5µm. **J)** Cartoon illustrating localization of sLro1 at Fld1 sites in the absence of Pet10.

### Pex30 is required downstream of Fld1, Nem1, and Yft2 to recruit the TAG-synthase

We wondered whether other proteins involved in LD biogenesis are important in organizing ER subdomains to facilitate droplet assembly. Pex30 is a reticulon-like ER membrane shaping protein (Joshi et al., 2016), that localizes to Fld1-containing ER subdomains (Joshi et al., 2018; Wang et al., 2018). In cells lacking Fld1, Pex30 mislocalizes to one large punctum per cell (Joshi et al., 2018; Wang et al., 2018). Since we find that Fld1, and Nem1 define ER sites of LD biogenesis, we wondered whether Pex30 plays a role upstream or downstream of Fld1 and Nem1 in organizing these ER subdomains. Hence, we tagged Fld1-mCherry, and Nem1-mCherry in *pex30*Δ mutants expressing GFP-FFAT-sLro1. Unlike in *fld1*Δ cells where Pex30 is completely mislocalized to one punctum per cell (Joshi et al., 2018; Wang et al., 2018), in cells missing Pex30, Fld1 or Nem1 still localized to discrete spots in the ER (Fig. 7D, E). This suggests that localization of Fld1 and Nem1 proteins is independent of Pex30. However, in the absence of Pex30, these subdomains failed to recruit GFP-FFAT-sLro1 (Fig. 7D, E). This indicates that Pex30 plays an indispensable role downstream of Fld1 and Nem1 in organizing ER subdomains for LD biogenesis (Fig. 7G).

To determine whether Yft2 functions upstream or downstream of Pex30, we generated *pex30*Δ cells expressing Fld1-mCherry or Nem1-mCherry, and co-expressing Yft2-sf-GFP. When these cells were grown in OA media to stimulate production of LDs, Yft2 showed punctate distribution and mostly colocalized with Fld1 or Nem1 (Fig. 7F). This suggests that localization of Yft2 at Fld1 or Nem1 sites is independent of Pex30 and implies that Pex30 is required downstream of Fld1, Nem1, and Yft2 in organizing ER subdomains for the recruitment of GFP-FFAT-sLro1 at LD biogenesis sites (Fig. 7H).

### Role of yeast PLIN proteins in organizing ER sites of LD biogenesis

Next, we tested whether Pet10, a yeast homolog of mammalian perilipins is required for the recruitment of GFP-FFAT-sLro1 at Fld1 or Nem1 sites (Gao et al., 2017). Perilipins are scaffolding proteins that stabilize LDs and play an important role in regulating storage and turnover of neutral lipids (Brasaemle, 2007). Yeast cells missing Pet10 display delayed LD biogenesis, decreased TAG accumulation in the ER, and fusogenic LDs (Gao et al., 2017). *pet10*Δ mutant cells expressing Fld1-mCherry or Nem1-mCherry and co-expressing GFP-FFAT-sLro1 showed no aberrant localization of Fld1 or Nem1, and did not impair the recruitment of GFP-FFAT-sLro1 at Fld1 sites (Fig. 7I, Fig S4H). This suggests that Pet10 acts downstream of Fld1, and Nem1 and that the yeast perilipin homologue is not required for the recruitment of the TAG-synthase at Fld1 or Nem1 sites (Fig. 7J).

### Loss of seipin results in the formation of ectopic LDs

Our results suggest that Fld1 and Nem1 define ER subdomains for the recruitment of the TAG-synthase and the LD biogenesis factors, Yft2, and Pex30. Thus, we tested if the LD marker, Erg6 would colocalize with Fld1 or Nem1 when cells are stimulated to produce LDs. Therefore, we generated 3ΔKO cells expressing genomic Fld1-mCherry or Nem1-mCherry and co-expressing Erg6-GFP from a plasmid. Prior to induction, Fld1 and Nem1 showed punctate localization in cells devoid of neutral lipids, and Erg6 was uniformly distributed over the ER membrane (Fig. 8A, B). Upon galactose induction, Erg6-GFP rapidly colocalized to ∼100% with Fld1 or Nem1 spots (Fig. 8A-C). Upon continued TAG synthesis, mature LDs remained associated with Fld1 or Nem1 sites (Fig. 8A, B), which is in agreement with previous findings showing that Fld1 and Nem1 localize to ER-LD contact sites (Adeyo et al., 2011; Grippa et al., 2015; Wolinski et al., 2015).

**Figure 8.**
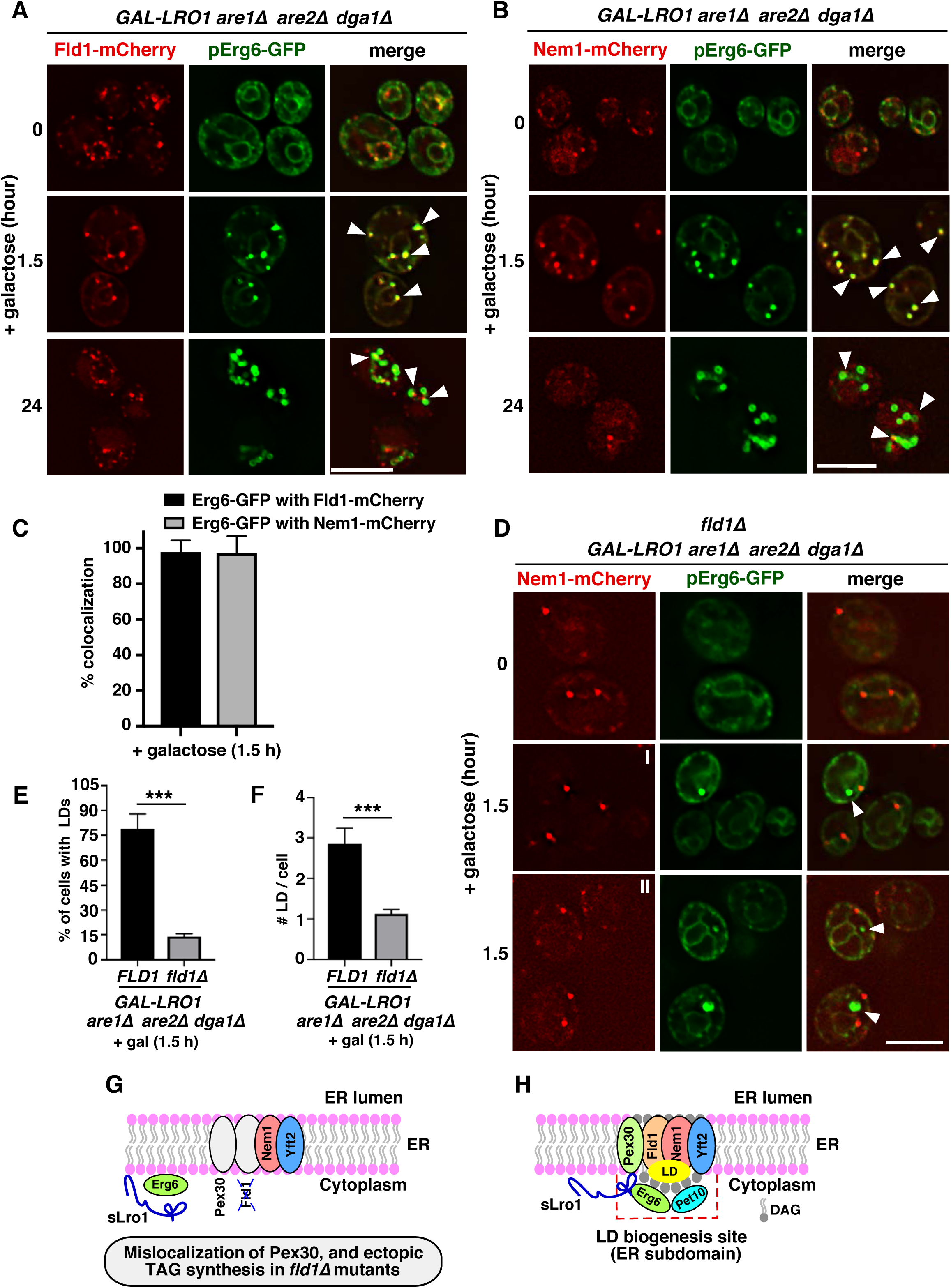
Absence of Fld1 from LD biogenesis sites result in ectopic droplet formation. **A, B)** LDs form at Fld1, and Nem1 marked ER sites. 3ΔKO expressing Fld1-mCherry (A) or Nem1-mCherry (B) and co-expressing Erg6-GFP were transferred to galactose containing media for the indicated time. White arrowheads denote colocalization between Erg6 with Fld1 (A) or Nem1 (B). **C)** Quantification of colocalization of Erg6 with Fld1 and Nem1. Data represent mean ± s.d., n > 50 cells. **D)** LDs fail to form at Nem1 sites in cells lacking Fld1. *fld1*Δ mutant cells (*fld1*Δ 3ΔKO) expressing Nem1-mCherry, and co-expressing Erg6-GFP, were grown as described in A. White arrowheads indicate Erg6 punctae that do not colocalize with Nem1. Scale bars: 5µm. **E, F)** Quantification of cells with at least 1 LD (E), or number of LDs per cell (F) for images shown in D. Data represent mean ± s.d., n > 50 cells; \*\*\**p* < 0.0005. **G)** Cartoon depicting lack of Fld1 from LD biogenesis sites does not affect the localization of Nem1 and Yft2, but results in mislocalization of Pex30 and failure to recruit the TAG-synthase. **H)** Cartoon showing that ER subdomains containing Fld1, Nem1, Yft2, and Pex30 define sites for the localization of the TAG-synthase and droplet formation. Erg6 and Pet10 bind to newly formed LDs from the cytosolic side.

**Figure 9.**
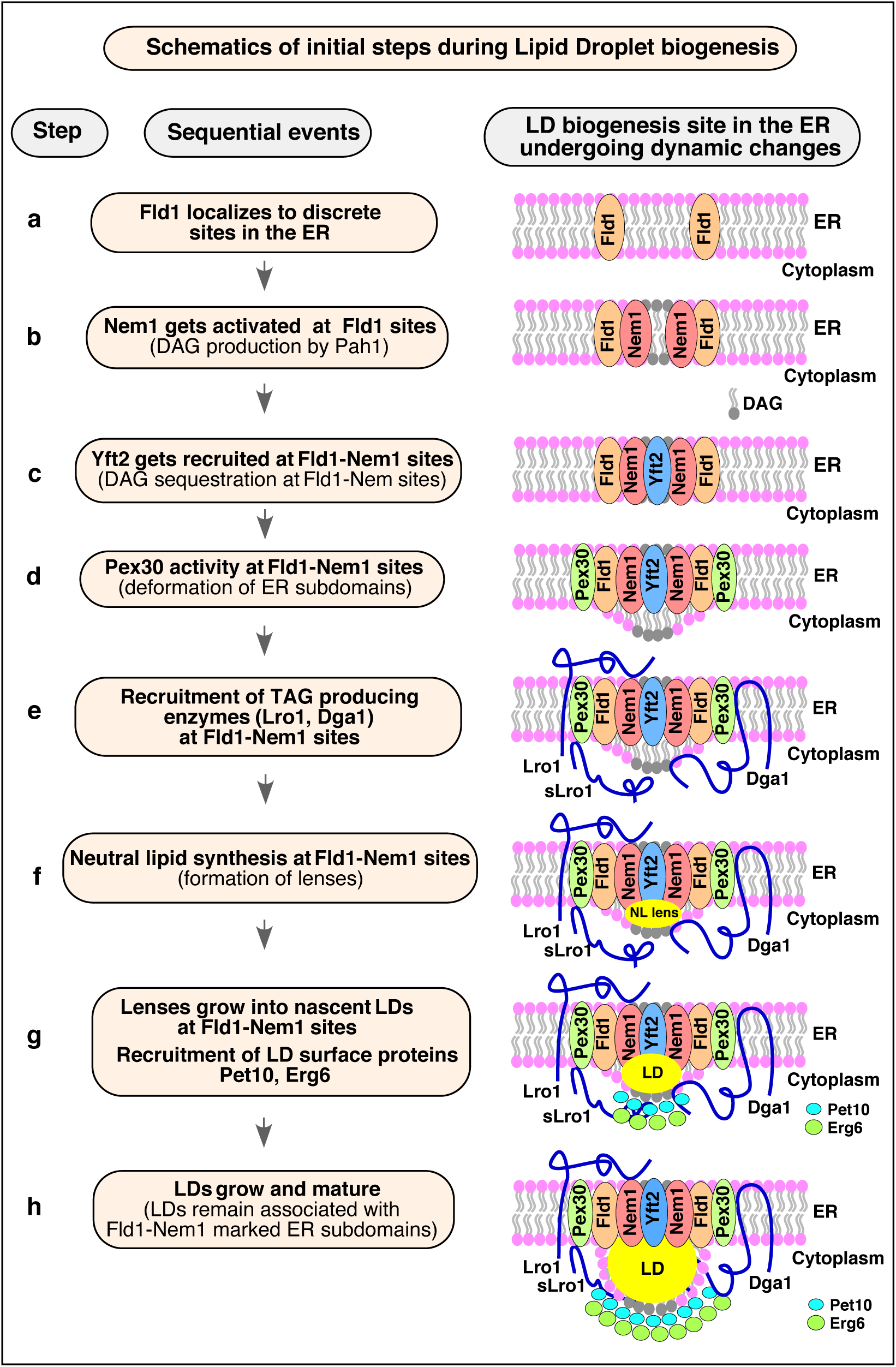
Model of LD formation at discrete ER subdomains defined by Fld1 and Nem1. Cartoon showing the order of events occurring at pre-defined ER subdomains for efficient LD formation. Fld1 and Nem1 marked ER subdomains undergo sequential enrichment with LD biogenesis factors thereby promoting localized TAG production and droplet assembly.

The fact that *fld1*Δ does not impair the localization of Nem1 into discrete ER sites, made us wonder whether Erg6 would still localize to Nem1 sites in the absence of Fld1. To examine this possibility, we used an inducible mutant lacking Fld1 (*fld1*Δ 3ΔKO) expressing Nem1-mCherry and Erg6-GFP. In cells lacking LDs, Erg6 showed uniform ER localization whereas Nem1 showed punctate ER distribution (Fig. 8D). Upon galactose induction, however, most of the cells failed to produce LDs as evidenced by few Erg6 spots (Fig. 8D). Only ∼10% of cells showed a single Erg6 punctum per cell, which did not colocalize with Nem1 (Fig. 8D-F). Deletion of Fld1 does not impair the rate of TAG synthesis, however, TAG accumulates throughout the ER membrane (Cartwright et al., 2015). In agreement with this, we find that in *fld1*Δ cells, induction of TAG synthesis resulted in accumulation of BODIPY throughout the ER, with a few BODIPY puncta that did not colocalize with Nem1 (Fig. S4I). This suggests that Fld1 activity at Nem1 sites plays a crucial role in the biogenesis of LDs, as in the absence of Fld1, TAG synthesis occurred throughout the ER. Fld1 is required for the proper localization of Pex30 (Joshi et al., 2018; Wang et al., 2018), though lack of Fld1 has no effect on the localization of Nem1 and Yft2 (Fig. 6E). Absence of Fld1 or Pex30 results in failure to recruit the TAG-synthase at ER subdomains containing Nem1 and Yft2, leading to ectopic TAG synthesis (Fig. 8G). Overall, these results indicate that LD biogenesis proteins, Fld1, Nem1, Yft2, Pex30, Pet10, Erg6, and Lro1 cooperate to define ER subdomains that are likely enriched in certain lipids to facilitate localized TAG synthesis and droplet formation (Fig. 8H).

### Induction of TAG-synthase Dga1 results in LD formation at Fld1 and Nem1 sites

Next, we asked if induction of the second TAG-synthase in yeast, Dga1, would also result in LD formation at Fld1 and Nem1 ER sites. Dga1, has a hairpin topology having both N- and C-termini exposed toward the cytosol, and catalyzes TAG formation from the cytosolic leaflet of the ER using membrane-embedded DAG, and cytosolic acyl-CoA as substrates (Oelkers et al., 2002; Sorger and Daum, 2002). Therefore, we generated 4ΔKO cells expressing Fld1-mCherry (Fig. S5A, C) or Nem1-mCherry (Fig. S5B, D) and co-expressing Dga1 from a galactose inducible promoter. Upon induction, BODIPY marked LDs rapidly appeared at Fld1 and Nem1 sites (Fig. S5A, B) and these sites were also marked by Erg6-GFP (Fig. S5C, D). Next, we investigated whether the TAG-synthase Dga1 also became enriched at Fld1 and Nem1 sites. Therefore, 4ΔKO cells expressing Fld1-GFP (Fig. S5E), or Nem1-GFP (Fig. S5F) together with Dga1-mCherry from a galactose promoter were analyzed. Prior to induction, Fld1 and Nem1 showed punctate localization (Fig. S5E, F). Upon galactose induction, these punctae rapidly became stained by Dga1-mCherry (Fig. S5E, F). These data suggest that not only Lro1, but also Dga1 localize to Fld1 and Nem1 marked ER subdomains during induction of LD biogenesis to promote localized TAG production and droplet assembly (Fig. S5G).

## DISCUSSION

What determines the sites of LD biogenesis in the ER is not well understood. In this study, we demonstrate that biogenesis of LDs occurs at pre-defined discrete ER subdomains marked by Fld1 and Nem1. The localization of both Fld1 and Nem1 at these specialized ER sites is independent of neutral lipid synthesis, or the presence of LDs. Fld1 and Nem1 localize independent of each other but both are required to create functional sites of LD biogenesis. These Fld1 and Nem1 marked ER sites become enriched with LD biogenesis proteins, including Yft2, Pex30, Pet10, Erg6, and Lro1. Based on these data, we propose a model for the stepwise assembly of factors for initiating LD biogenesis (Fig. 10).

In cells devoid of neutral lipids, Fld1 and Nem1 localize to discrete ER sites (Fig. 10a, b). We do not yet know the determinant for this punctate localization. It is possible that specific proteins, lipids, or possibly a combination of both facilitates the recruitment of Fld1 and Nem1 into discrete spots in the ER in cells lacking LDs. This should be investigated in further studies. Almost all of the Nem1 punctae appear to be stable and immobilized in the ER, whereas some of the Fld1 punctae move along the ER while others are stable. It is likely that only the sites at which both Fld1 and Nem1 colocalize (Fld1-Nem1 sites) become functional for droplet assembly. Consistent with this a new study identified seipin in complex with LDAF1 (lipid droplet assembly factor 1) that copurified with TAG, and defined sites of LD biogenesis (Chung et al., 2019). Moreover, mislocalization of seipin within the ER network to the nuclear envelop results in formation of LDs at these ectopic sites (Salo et al., 2019).

Fld1/Nem1 sites require the activity of Yft2, and Pex30, to ensure proper recruitment of the TAG-synthase, indicating that all four proteins (Fld1/Nem1/Yft2/Pex30) might work together in establishing ER subdomains permissive for localized TAG synthesis. Yft2 gets recruited to Fld1/Nem1 sites (Fig. 10c), possibly through direct binding of DAG and/or TAG (Gross et al., 2011). We have recently shown that Yft2 colocalizes with Fld1, Nem1, and the ER-DAG sensor (Choudhary et al., 2018). Pex30, acts downstream of Fld1, Nem1, and Yft2, to localize to LD biogenesis sites (Fig. 10d). The membrane shaping property of Pex30 at these sites may facilitate the deformation of the ER bilayer to accommodate DAG and/or TAG. Indeed, Pex30 at LD biogenesis sites colocalizes with Fld1, Nem1, and the ER-DAG sensor (Joshi et al., 2018; Wang et al., 2018).

Next, the TAG-synthase, Lro1/Dga1 localizes to Fld1/Nem1/Yft2/Pex30 sites resulting in the conversion of DAG to TAG and hence formation of lenses that grow into nascent LDs (Fig. 10e, f, g). The fact that the sLro1 goes to discrete ER sites suggests that its localization is driven by lipids since in the native Lro1 this portion of sLro1 is in the ER lumen and thus would not be available to specific protein-protein interactions in the cytosolic leaflet of the ER membrane. This hypothesis is supported by three findings: first, native Lro1, and sLro1 colocalize with the ER-DAG sensor (Fig. 4C), second, purified sLro1 binds to DAG *in vitro* (Fig. 4F), third, the mislocalizatio of Lro1 in *nem1*Δ cells is rescued by the expression of a hyperactive Pah1 (Pah1-7P) (Fig. 5G). In addition, proper recruitment of Lro1 requires the enzymatic activity of Lro1 as a catalytic dead point mutant of sLro1 (H618A) that still binds DAG *in vitro*, fails to enrich at Fld1 sites *in vivo* (Fig. 4G, H).

Nascent LDs are then recognized by cytosolic proteins such as the perilipin homolog Pet10, and Erg6, and the LD monolayer surface becomes stabilized, facilitating LD growth toward the cytosol (Fig. 10g, h). Consistent with this proposition, artificial LD embedded into giant unilamellar vesicle emerge toward the side having greater coverage with proteins and phospholipids, which reduce surface tension of the monolayer (Chorlay et al., 2019). Continued synthesis of TAG at Fld1/Nem1 sites results in maturation of LDs, which remain associated with the ER membrane at Fld1/Nem1 sites (Fig. 10h). In conclusion, our findings support the hypothesis that biogenesis of LDs is a well-orchestrated process that occurs at Fld1/Nem1/Yft2/Pex30 marked discrete ER subdomains.

## Acknowledgements

We thank Dr. Paolo Ronchi and the Electron Microscopy Core Facility of the European Molecular Biology Laboratory Heidelberg for expert technical assistance in CLEM imaging. This work was supported by the Swiss National Science Foundation (Grant # 31003A_17303), the Novartis Foundation for medical-biological Research (Grant # 19B140), and the Intramural Research Program of the National Institute of Diabetes and Digestive and Kidney Diseases. Vineet Choudhary is supported by Marie Curie fellowship from the European Commission (Grant # 747536, LD_Biogenesis).

## Author Contributions

Vineet Choudhary conceived and performed experiments, analyzed the data, and wrote the paper. Ola El Atab expressed and purified proteins and performed *in vitro* DAG binding assay. Giulia Mizzon performed EM experiments. William A Prinz conceived experiments, analyzed the data, and edited the paper. Roger Schneiter conceived experiments, analyzed the data, and wrote the paper.

## Declaration of Interests

The authrs declare no competing interests.

## Materials and Methods

### Yeast strains and plasmids

Yeast strains and plasmids used in this study are listed in Supplementary Table S1. Double- and triple-deletion mutants were generated either by mating or by PCR-based targeted homologous recombination to replace the ORF of the gene of interest with a deletion cassette followed by a marker rescue strategy (Gueldener et al., 2002). Chromosomally C-terminal tagged versions of reporter proteins (Erg6-mCherry, Nem1-mCherry, Fld1-mCherry, Fld1-GFP, Nem1-GFP,) were constructed using PCR-based homologous recombination (Longtine et al., 1998). The cassette yEmCherry-*HIS5MX6* was obtained from J. Nunnari (University of California, Davis, CA). To construct the plasmid pGAL1-GFP-Lro1 (amino acids 1-661), or pGAL1-GFP-sLro1 (amino acids 98-661), corresponding regions of *LRO1* were amplified by PCR from *S. cerevisiae* genomic DNA and inserted into the plasmid pRS426 thereby expressing GFP fused to the N-terminus of Lro1, or sLro1 under the *GAL1* promoter. For plasmid pGAL1-GFP-FFAT, GFP was fused to a FFAT motif and inserted into pRS426 under GAL1 promoter. To create the plasmid pGAL1-MBP-GFP-FFAT-sLro1, the sLro1 (98-661) sequence was fused to FFAT, GFP and the purification tag maltose binding protein (MBP), and inserted into pRS426. To construct pGAL1-MBP-Scarlet-FFAT-sLro1, sequence encoding Scarlet tag was amplified from lab plasmid collection, and fused with FFAT, sLro1 (98-661) and MBP tag and inserted into pRS426. PCR mediated site directed mutagenesis was performed to generate sLro1^H618A^ and cloned as a fusion with MBP-Scarlet-FFAT-sLro1^H618A^ under GAL1 promoter into pRS426 vector. To generate pGAL1-ssKar2-GFP-sLro1, the sequence corresponding to sLro1 (98-661) was fused to GFP and the signal sequence of Kar2 (amino acids 1-42), and cloned into pRS426. For plasmid pGAL-Sec63(1-250)-GFP-sLro1, the sequence corresponding to 1-250 amino acids encoding TMDs (1-3) were fused with GFP, and sLro1 (98-661) and inserted into pRS426 vector. We used plasmids expressing either mCherry-HDEL, RFP-HDEL, or GFP fused to Sec63 to label the ER. pADH-Fld1-mCherry was generated by amplifying the *FLD1* gene from *S. cerevisiae* genomic DNA and inserting it into pRS415. The plasmids encoding Yft2-sf-GFP, and the ER-DAG sensor (pADH-ER-DAG sensor) have been described previously (Choudhary et al., 2018; Choudhary et al., 2015). Plasmid expressing hyperactive allele of Pha1 (Pah1-7P) was obtained from S. Siniossoglou (University of Cambridge, UK). All constructs were verified by sequencing.

### Media and growth conditions

Cells were grown at 30°C in YPD medium (1% Bacto yeast extract, 2% Bacto Peptone, 2% glucose) unless otherwise indicated. Cells containing plasmids were grown in synthetic complete (SC) media containing 6.7 g/l yeast nitrogen base without amino acids (United States Biologicals), an amino acid mix (United States Biologicals), containing either 2% glucose, galactose or raffinose. To grow cells in presence of fatty acids for induction of LD formation, SC media were supplemented with 0.1% oleic acid (Sigma Aldrich) in the presence of 1% Brij-58 (Sigma Aldrich).

### Fluorescence microscopy

For Fig. 1E, 1F, and supplementary Fig. S1B, C, cells were visualized and imaged live in culture media using an Olympus BX61 microscope, a UPlanApox100/1.35 lens, a QImaging Retiga EX camera, and processed using IVision software (version v 4.0.5). Images being directly compared were obtained using identical microscope settings. For Fig. 2 – 8, and supplementary Fig. S1F, S2 – 5, live cell imaging was performed using a DeltaVision Elite imaging system (GE Healthcare, Pittsburg, PA). The DeltaVision Elite is comprised of a wide field inverted epifluorescence microscope (1×71, Olympus), equipped with a charge-coupled device (CCD) camera (CoolSNAP HQ^2^, Photometrics, Tuscon, AZ). Images were acquired using a U PLAN S-APO 100×1.42 NA oil immersion objective (Olympus) and a GFP or mCherry filter set. Stacks of seven images with a step size of 0.5µm were deconvolved for 10 iterations using the conserved ratio method in softWoRx (Applied Precision). Single plane images are displayed. Time-lapse images were acquired every 1 s for 15 s with an exposure of 500 ms for Fig. S3B and every 1 s for 60 s with an exposure of 500 ms for Fig. S3C. Brightness and contrast were adjusted using Adobe Photoshop CC 2020 (Adobe Systems). Colocalization between GFP and mCherry marked punctae in acquired images was scored manually. Values were recorded in Microsoft Excel and analyzed in Prism 8.0 (GraphPad Software).

### BODIPY staining to visualize lipid droplets in yeast

To visualize LDs in yeast, cells expressing Erg6-mCherry, were grown in YPD to early stationary phase and stained with 0.5 µg/ml BODIPY 493/503 (Invitrogen) for 5 min at RT. After washing once with 1x PBS, cells were visualized live using the GFP or mCherry filter set of the DeltaVision Elite (GE Healthcare, Pittsburg, PA).

### Labeling with [^3^H]palmitic acid and quantification of lipids

To quantify neutral lipids and phospholipids, cells were grown in SC raffinose media overnight. Next day cells were diluted to 1 OD_600_ per ml and grown in SC galactose media containing 10 µCi/ml [^3^H]palmitic acid (10 mCi/ml; American Radiolabled Chemicals, Inc., St Louis, MO). Cells were incubated with shaking at 30°C. 10 OD_600_ units of cells were harvested at 10-, 30- 60-, 90-, 120-min (Fig S4F), at 10-min, 2 h, 4 h, 21 h (Fig. 1C), and at 21 h (Fig. 1D, 4D, S1G). Cells were lysed using a Precellys 24 homogenizier, lipids were extracted as described (Jacquier et al., 2011) and equal aliquots were brought to dryness. To quantify TAG and SE, lipids were spotted onto silica gel 60 thin layer chromatography (TLC) plates (Merck, Darmstadt, Germany) and developed with petroleum ether/diethylether/acetic acid (70:30:2; per vol). Glycerophospholipids were separated by TLC as described (Vaden et al., 2005). TLC plates were exposed to a tritium-sensitive screen, visualized and quantified using a phosphorimager (GE Healthcare). Brightness and contrast of the TIFF images were adjusted using Image J (NIH).

### Western blot analysis

Cells were grown over night in SC raffinose media, diluted to 1 OD_600_ ml in SC galactose media and cells corresponding to 3 OD_600_ units were harvested at 0-, 2-, 4-, 6-, 21-h time points. Samples were extracted by NaOH, and proteins were precipitated using trichloroacetic acid (TCA), and resuspended into sample buffer (Horvath and Riezman, 1994). Proteins were separated using SDS-PAGE and detected using a monoclonal antibody against GFP (Roche Diagnostics, Rotkreuz, Switzerland, dilution 1:2000), or Kar2 (Santa Cruz Biotechnology, Dallas, TX, dilution 1:5000) and using horseradish peroxidase (HRP)-conjugated secondary antibodies (Santa Cruz Biotechnology, Dallas, TX, #2302, dilution 1:10000). Western blot experiments were performed at least twice with similar results.

### Quantification and statistical analysis

Unless otherwise stated, data were obtained from at least three independent experiments. The images presented are representative of the results obtained. Quantitative results are expressed as mean ± standard deviation. Statistical analysis was performed with unpaired, two-tailed Students’s t-tests using GraphPad Prism 8 software (GraphPad, La Jolla, CA). p-values < 0.05 were considered significant.

### Correlative light and electron microscopy (CLEM)

For CLEM investigations, yeast cells were concentrated to a paste by vacuum-filtration and subsequently cryoimmobilised with an HPM010 high pressure freezing machine (Bal-Tec, Los Angeles, CA, USA). Automated freeze-substitution in 0.1% uranyl acetate in acetone over a 72 h period and embedding in Lowicryl HM20 resin were performed using an EM AFS2 (Leica Microsystems) as previously described (Kukulski et al., 2011; Thaller et al., 2019). Blocks were polymerized under UV light, cut in 300 nm thick sections using an ultramicrotome (EM UC7, Leica Microsystems) and sections were picked up on carbon coated 200 mesh grids (S160, Plano). 100 nm or 50 nm TetraSpeck fluorescent microspheres (Lifetechnologies, Carlsbad, CA, USA) were adsorbed to the grid for the subsequent fiducial-based correlation between light and electron microscopy images. The fluorescence microscopy (FM) imaging of the sections was carried out as previously detailed (Curwin et al., 2016; Kukulski et al., 2011) using a widefield fluorescence microscope (Olympus IX81) equipped with an Olympus PlanApo 60X 1.40 NA oil immersion objective. In addition to the specific channel of the fluorescent protein(s) of interest another channel was acquired in order to distinguish the specific signal from the Tetraspecks. Grids were then post-stained for 5 min with 1% uranyl acetate and 3 min with Reynolds lead citrate. Tilt series were acquired with a FEI Tecnai F30 electron microscope run at 300kV and controlled by Serial EM (Mastronarde, 2005). Fluorescence microscopy maps were loaded using a point-based registration to low mag tiled acquisitions (2,300x) of the corresponding grid squares. High-magnification tilt series (15,500x) were then acquired at the selected fluorescent spot locations, using a Serial EM script (Schorb et al., 2019). Tomograms were then reconstructed using the IMOD software package (Kremer et al., 1996). For samples with 50 nm Tetraspeck beads an intermediate acquisition at 4,700x was necessary to perform the alignment, as the beads were not visible in the lowest magnification. CLEM overlays were performed with a fiducial-based correlation using the plugin eC-CLEM in Icy (Paul-Gilloteaux et al., 2017) as previously described (Curwin et al., 2016). Tomograms were manually segmented using IMOD (Kremer et al., 1996).

### Expression and purification of wild-type and mutant versions of sLro1

DNA encoding wild-type and mutant versions of sLro1 were cloned into the BamHI and XhoI sites of pET22b vector (Novagen, Merck, Darmstadt, Germany), which contains a PelB signal sequence to direct the secretion of the expressed protein into the periplasmic space. Plasmids were transformed into *Escherichia coli* BL21 and the proteins were expressed with a C-terminal polyhistidine-tag after overnight lactose induction and growth of bacteria at 24°C. Cells were collected, lysed and incubated with nickel-nitrilotriacetic acid beads (Qiagen, Hilden, Germany) as per the manufacturer instructions. The beads were washed, loaded onto a Ni^2+^-NTA column (Qiagen) and proteins were eluted in 60 mM NaH_2_PO_4_, 60 mM NaCl and 300 mM imidazole, pH 8.0. The eluted protein was applied to Zeba™ spin desalting columns (Thermo scientific) and the buffer was exchanged to 60 mM NaH_2_PO_4_, 10 mM NaCl, pH 8.0. Protein concentration was determined by Lowry assay using folin reagent and BSA standard.

### Microscale Thermophoresis (MST)

Microscale thermophoresis experiments were performed using a Monolith NT.115 from Nanotemper Technologies (Munich, Germany) to assess the binding of sLro1 and sLro1-H618A to short chain DAG (C8:0 DAG, catalog number: 800800, Avanti Polar Lipids, USA). Proteins were labeled using the RED-tris-NTA His tag protein labeling kit (Nanotemper Technologies). Labelled protein was added to a serial dilution of unlabeled DAG, prepared in binding buffer (20 mM Tris pH 7.5, 30 mM NaCl, 0.05% Triton X-100). Samples were loaded into MST standard capillaries, and MST measurements were performed using 80% power setting. The dissociation constant *Kd* was obtained by plotting the normalized fluorescence Fnorm against the logarithm of DAG concentrations. Experiments were performed in triplicates and data were fitted using the *Kd* model with the MO Affinity Analysis software.

### Data availability

The authors declare that the data supporting the findings of this study are contained with the paper and its supplementary information files.

## Supplementary Information

### Supplementary figure legends

**Figure S1.**
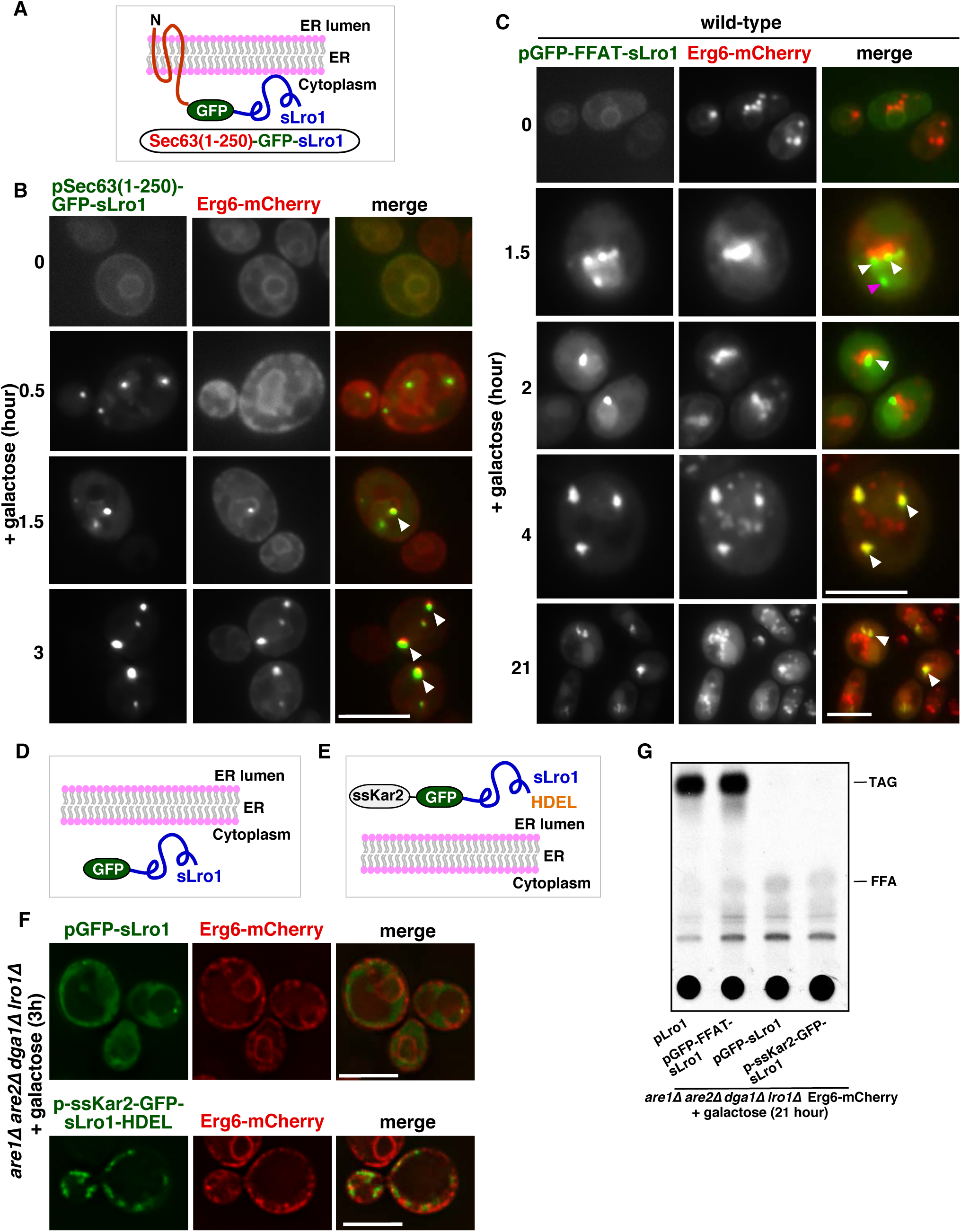
Data associated with Fig. 1. TAG-synthase sLro1 needs to be ER associated for its activity. **A, B)** sLro1 containing transmembrane domains from Sec63 localizes to ER subdomains. Design of the TAG-synthase chimera containing TMDs 1-3 from Sec63 (1-250aa), facing the active site to the cytoplasm {Sec63(1-250)-GFP-sLro1} (A). 4ΔKO cells expressing Erg6-mCherry and co-expressing Sec63(1-250)-GFP-sLro1 from a galactose inducible promoter were grown in raffinose and switched to galactose media for the indicated period of time (B). White arrowheads indicate colocalization of Sec63(1-250)-GFP-sLro1 punctae with Erg6-mCherry. Scale bar: 5µm **C)** GFP-FFAT-sLro1 localizes to LDs in wild-type cells. Wild-type cells expressing Erg6-mCherry and co-expressing GFP-FFAT-sLro1 from a galactose inducible promoter were grown as in B. White arrowheads indicate colocalization of GFP-FFAT-sLro1 punctae with Erg6-mCherry. The red arrowhead denotes a GFP-FFAT-sLro1 puncta not associated with Erg6-mCherry. Scale bars: 5µm. **D, E, F)** The TAG-synthase sLro1 needs to be anchored to the ER membrane for its activity. Design of the soluble fusion protein GFP-sLro1 localized in the cytosol (D), and ssKar2-GFP-sLro1-HDEL localized in the ER lumen (E). 4ΔKO cells expressing Erg6-mCherry, and co-expressing either GFP-sLro1 or ssKar2-GFP-sLro1-HDEL from a galactose inducible promoter. Cells were grown in raffinose and switched to galactose containing media for 3 h (F). Scale bars: 5µm. **G)** Non-membrane associated sLro1 fails to catalyze TAG synthesis. ΔKO cells expressing genomic Erg6-mCherry and co-expressing either native Lro1, GFP-FFAT-sLro1, GFP-sLro1, or ssKar2-GFP-sLro1-HDEL from a galactose inducible promoter were grown in galactose media containing [^3^H]-palmitic acid for 21 h. Lipids were extracted and separated by thin layer chromatography (TLC). TAG, triacylglycerol, FFA, free fatty acid.

**Figure S2.**
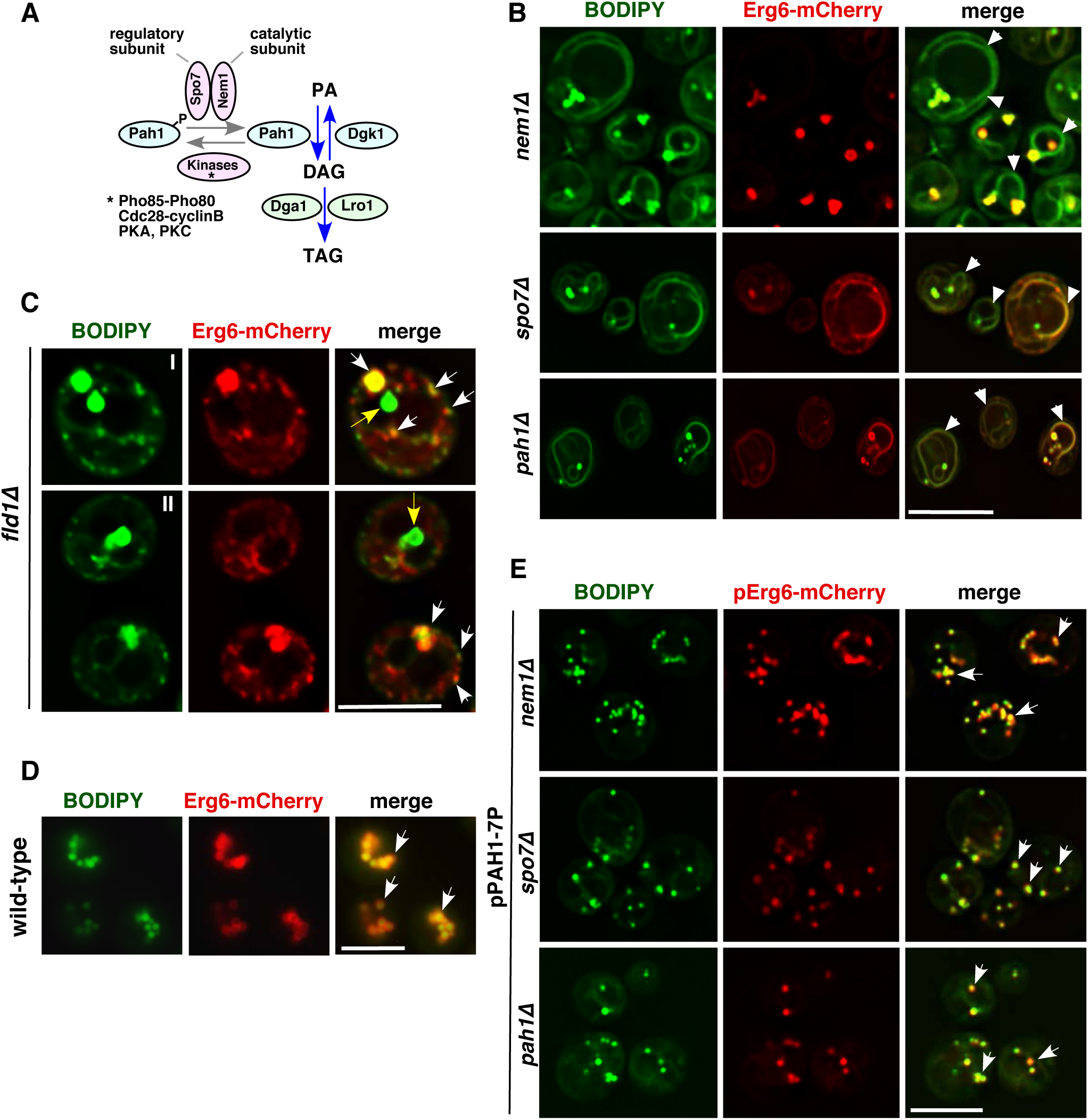
Data associated with Fig. 2. Cells lacking Nem1, Spo7, Pah1 or Fld1 accumulate neutral lipids in the ER and have aberrant droplets. **A)** Diagram depicting the regulation of triglyceride synthesis. The Nem1-Spo7 complex dephosphorylates and thereby activates the phosphatidate phosphatase, Pah1, resulting in formation of diacylglycerol (DAG) from phosphatidic acid (PA). DAG then serves as a substrate for the TAG synthesizing enzymes, Dga1 and Lro1. **B)** BODIPY accumulates in the ER of *nem1*Δ, *spo7*Δ, and *pah1*Δ cells. Yeast cells of the indicated genotype expressing genomic Erg6-mCherry were stained with BODIPY. White arrowheads denote BODIPY accumulation in the ER membrane. Scale bar: 5µm. **C)** Fld1 deleted cells accumulate numerous small and few supersized LDs. *fld1*Δ cells expressing genomic Erg6-mCherry were stained with BODIPY. White arrows denote colocalization between BODIPY and Erg6-mCherry. Yellow arrows denote BODIPY stained supersized puntae that do not colocalize with Erg6-mCherry. Scale bars: 5µm. **D)** Wild-type cells do not show BODIPY accumulation in the ER. BODIPY staining of wild-type cells expressing genomic Erg6-mCherry. White arrows denote colocalization between BODIPY and Erg6-mCherry. Scale bar: 5µm. **E)** Hyperactive Pah1 (Pah1-7P) rescues the LD biogenesis defect of *nem1*Δ, *spo7*Δ, and *pah1*Δ mutant cells. BODIPY staining of *nem1*Δ, *spo7*Δ, and *pah1*Δ mutants expressing Erg6-mCherry and a hyperactive allele of Pah1 (Pah1-7P). White arrows denote colocalization between BODIPY and Erg6-mCherry. Scale bar: 5µm.

**Figure S3.**
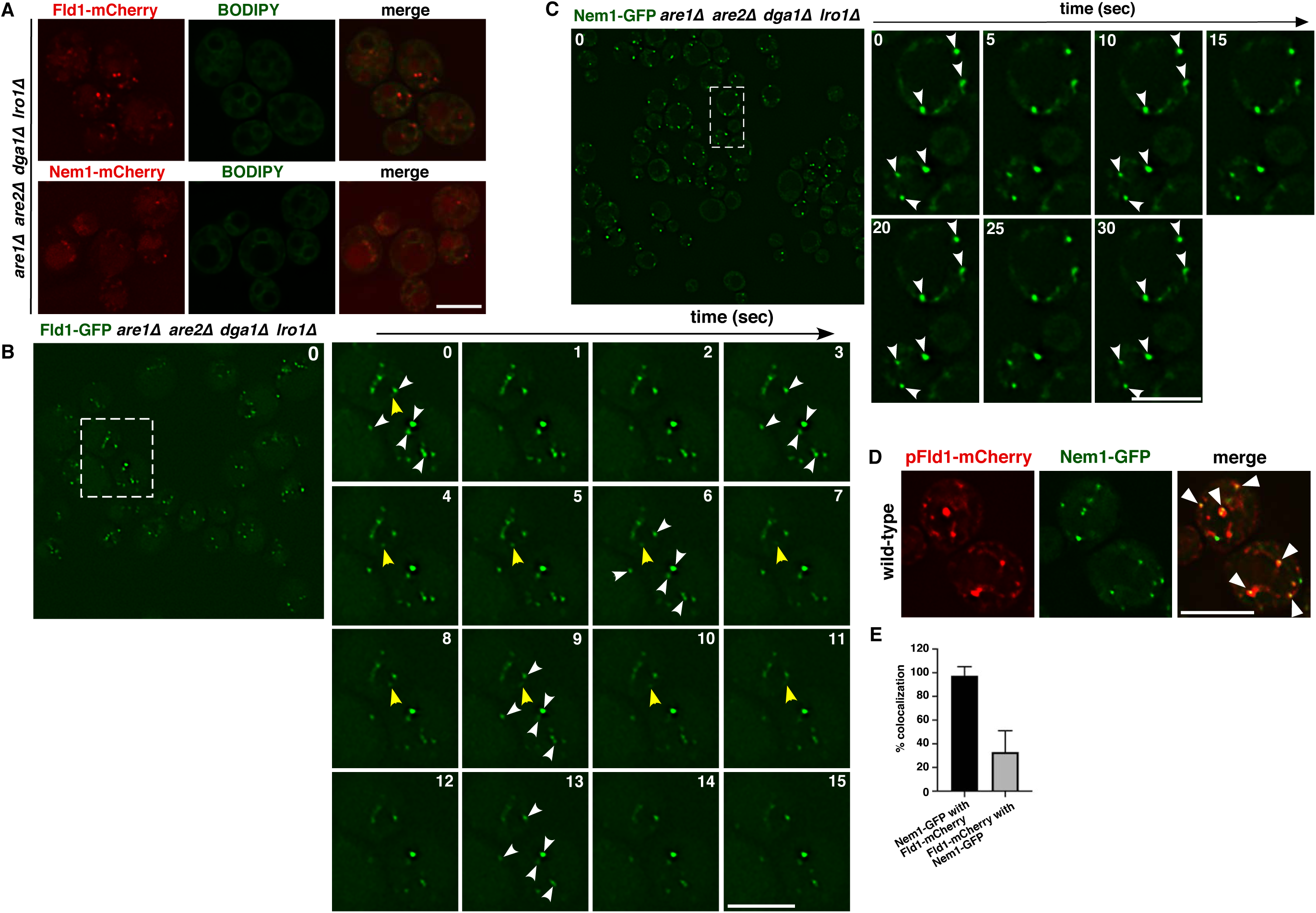
Data associated with Fig. 2 and 3. Fld1 and Nem1 localize to punctae in cells devoid of neutral lipids or LDs. **A)** Localization of Fld1 or Nem1 to ER foci is independent of neutral lipid synthesis or the presence of LDs. BODIPY staining of 4ΔKO cells expressing genomic Fld1-mCherry or Nem1-mCherry. Scale bar: 5µm. **B, C)** Time lapse images of Fld1-GFP and Nem1-GFP. 4ΔKO cells expressing genomic Fld1-GFP (B) or Nem1-GFP (C) were grown to early stationary phase and time lapse images were recorded. The boxed region is shown in higher magnification. Snapshots of images at the indicated time points are shown. White arrowheads indicate Fld1 or Nem1 punctae that stay immobile in the ER membrane. Yellow arrowheads denote rapidly moving Fld1 punctae. Scale bar: 5µm. **D)** Fld1 colocalizes with Nem1. Wild-type cells expressing Nem1-GFP and co-expressing Fld1-mCherry were imaged. White arrowheads denote colocalization between Fld1 and Nem1. Scale bar: 5µm. **E)** Quantification of colocalization between Nem1-GFP and Fld1-mCherry. Data represent mean ± s.d., n > 50 cells;

**Figure S4.**
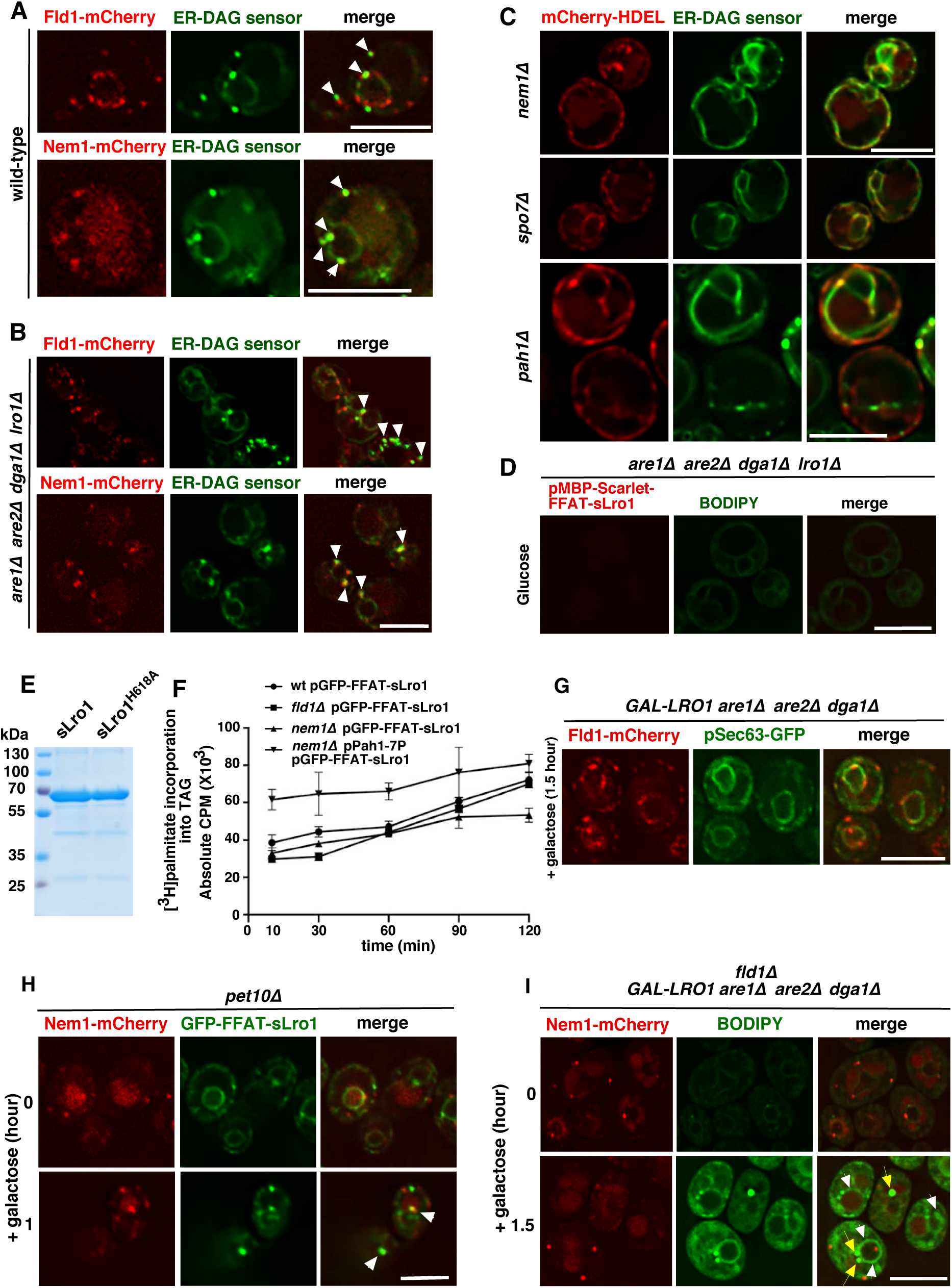
Data associated with Fig. 4, 5, 6, 7, 8. Enrichment of DAG at Fld1 and Nem1 ER subdomains. **A)** ER subdomains marked by Fld1 and Nem1 show enrichment of DAG. Wild-type cells expressing Fld1-mCherry, or Nem1-mCherry and co-expressing the GFP-tagged ER-DAG sensor were grown in SC media to early stationary phase and imaged live. White arrowheads indicate colocalization of the ER-DAG sensor with ER punctae marked with Fld1 or Nem1. Scale bars: 5µm. **B)** Fld1 and Nem1 marked ER subdomains show enrichment of DAG in LD-deficient cells. 4ΔKO cells expressing Fld1-mCherry or Nem1-mCherry and co-expressing the ER-DAG sensor were grown as in A. White arrowheads indicate colocalization of the ER-DAG sensor with punctae marked by Fld1 or Nem1. Scale bar: 5µm. **C)** Lack of Nem1, Spo7 or Pah1 results in uniform distribution of the ER-DAG sensor. *nem1*Δ, *spo7*Δ or *pah1*Δ mutant cells expressing the ER-DAG sensor were grown as in A. ER was visualized by mCherry-HDEL. Scale bar: 5µm. **D)** BODIPY staining of 4ΔKO cells expressing MBP-Scarlet-FFAT-sLro1. Cells were grown in glucose media to mid log phase. Scale bar: 5µm. **E)** Coomassie staining of an SDS-PAGE gel showing purified WT and mutant sLro1. **F)** Rate of TAG synthesis. Time dependent incorporation of [^3^H]palmitic acid into TAG in the indicated yeast mutant cells. Cells were labeled and collected at 10-, 30-, 60-, 90-, and 120-min time points. Data represent mean ± s.d. of three independent experiments. **G)** An ER membrane protein that is not involved in LD biogenesis does not show enrichment at Fld1 marked ER subdomains. The ER protein Sec63 does not become enriched at Fld1 sites during induction of LD biogenesis. *GAL-LRO1* 3ΔKO cells expressing Fld1-mCherry and co-expressing Sec63-GFP were grown in raffinose and switched to galactose containing media for 1.5 h. Scale bar: 5µm. **H)** Lack of Pet10 does not affect Nem1 localization and recruitment of TAG synthase. The recruitment of GFP-FFAT-sLro1 to Nem1 sites is not affected in *pet10*Δ mutant cells. *pet10*Δ mutant cells expressing Nem1-mCherry and co-expressing GFP-FFAT-sLro1 from a galactose inducible promoter were shifted to galactose media for the indicated period of time. Two examples of the 1 h time point are shown (I, II). Arrowheads denote colocalization of sLro1 with Nem1 foci. Scale bar: 5µm. **I)** Neutral lipids accumulate in the ER membrane in seipin mutant cells. *fld1*Δ mutant cells (*fld1*Δ *GAL-LRO1* 3ΔKO) expressing Nem1-mCherry were grown in raffinose and transferred to galactose containing media for the indicated period of time, stained with BODIPY and imaged. Two examples of the 1.5 h time point are shown (I, II). White arrowheads indicate BODIPY accumulation in the ER. Yellow arrows indicate BODIPY punctae that do not colocalize with Nem1 foci. Scale bar: 5µm.

**Figure S5.**
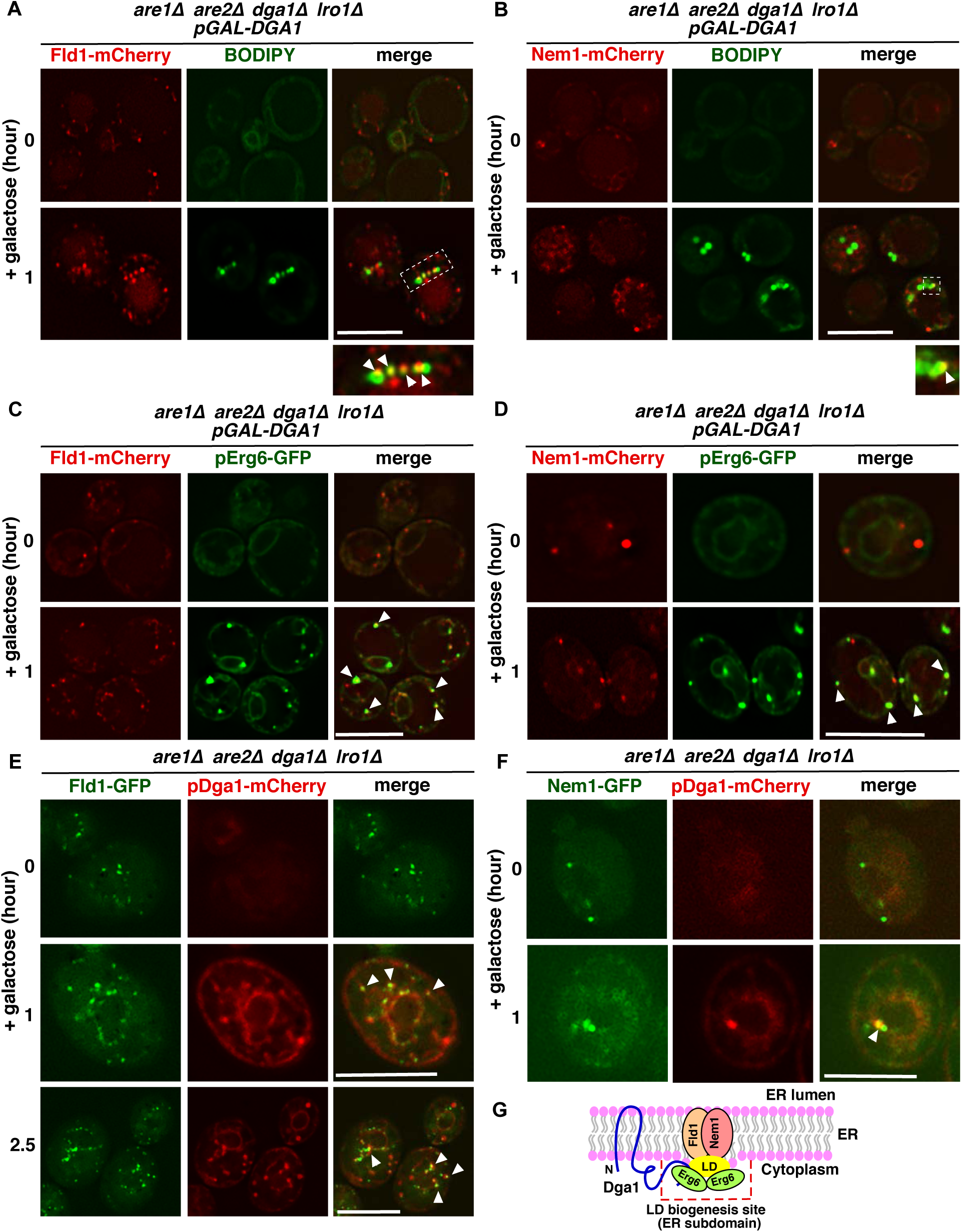
Induction of TAG-synthase Dga1 results in LD formation at Fld1 and Nem1 sites. **A, B)** Fld1 and Nem1 sites are stained with BODIPY upon TAG synthesis. 4ΔKO cells expressing Fld1-mCherry (A) or Nem1-mCherry (B), and co-expressing Dga1 from a galactose inducible promoter were switched to galactose media for 1 hour and stained with BODIPY. White arrowheads indicate colocalization between BODIPY and Fld1/Nem1 punctae. **C, D)** ER subdomains marked by Fld1 and Nem1 become enriched with LD marker protein. 4ΔKO cells expressing Fld1-mCherry (C) or Nem1-mCherry (D), Erg6-GFP, and co-expressing Dga1 from a galactose inducible promoter were grown as in A. White arrowheads indicate colocalization between Erg6 and Fld1/Nem1 punctae. **E, F)** Dga1 colocalizes with Fld1/Nem1 sites. 4ΔKO cells expressing Fld1-GFP (E) or Nem1-GFP(F) and co-expressing Dga1-mCherry from a galactose inducible promoter were switched to galactose for the indicate time. White arrowheads denote colocalization between Dga1-mCherry and Fld1-GFP/Nem1-GFP punctae. Scale bars: 5µm. **G)** Cartoon illustrating the recruitment of Dga1-mCherry to Fld1 and Nem1 ER sites, where droplets begin to form.

**Supplementary Table 1.**
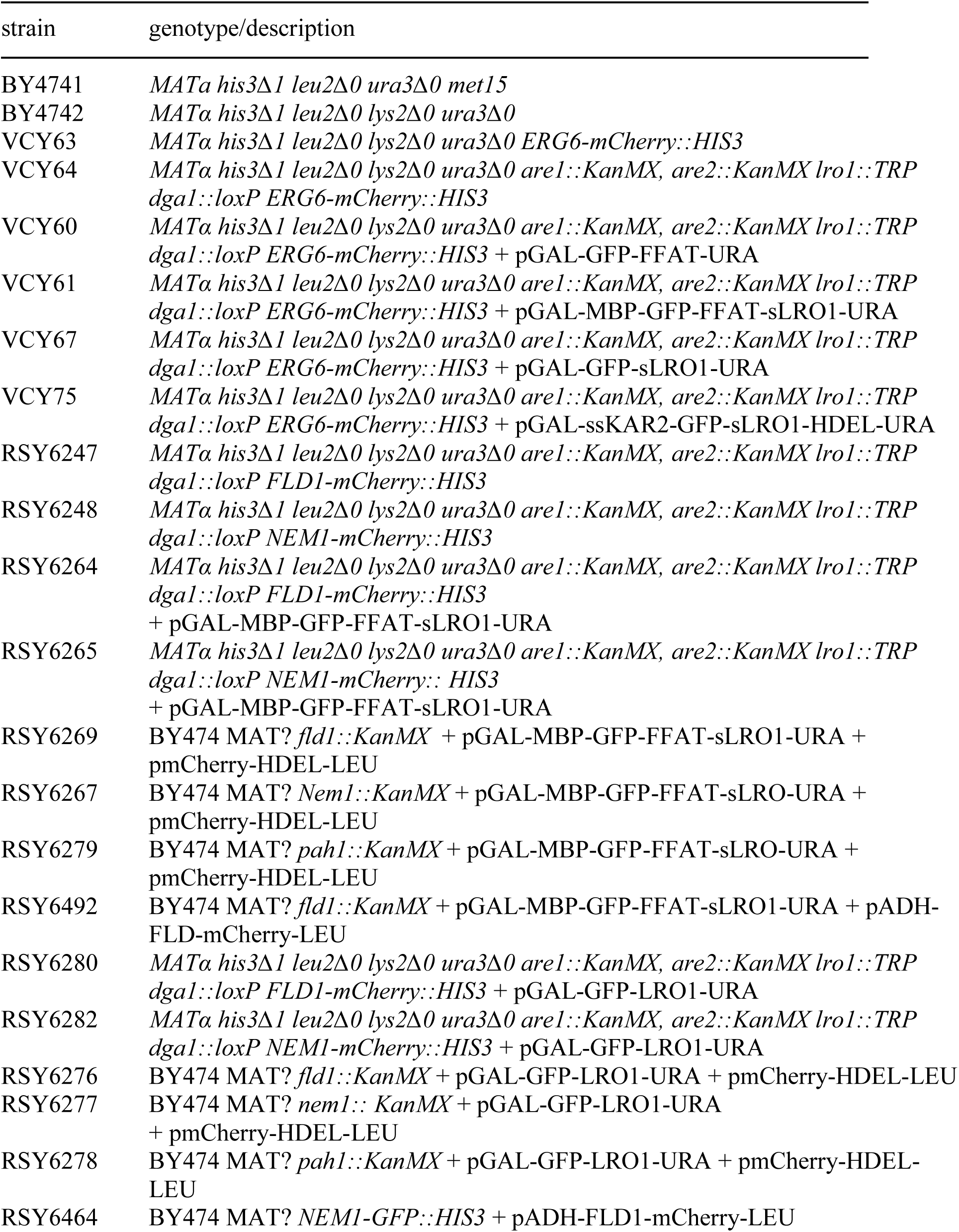
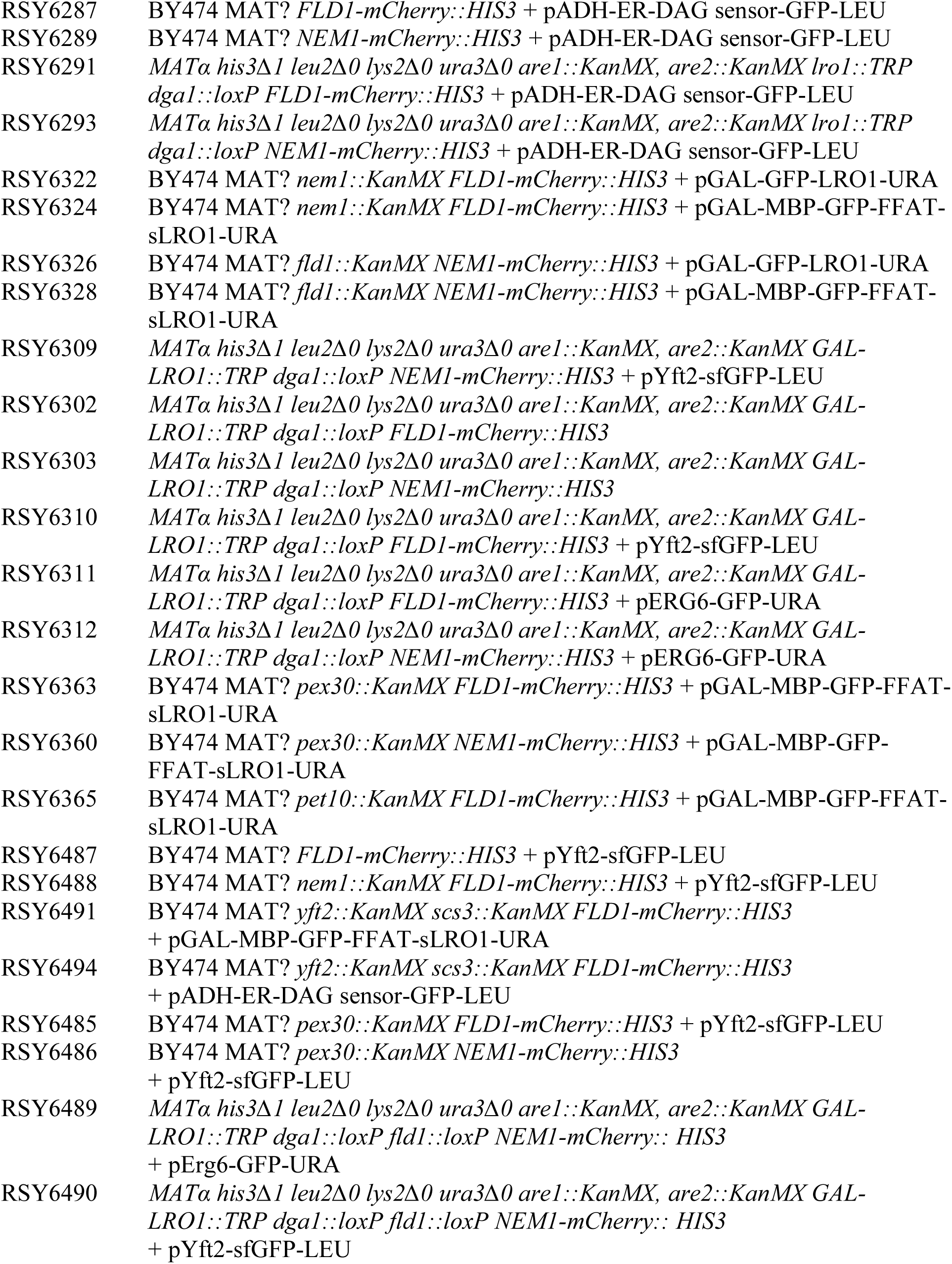
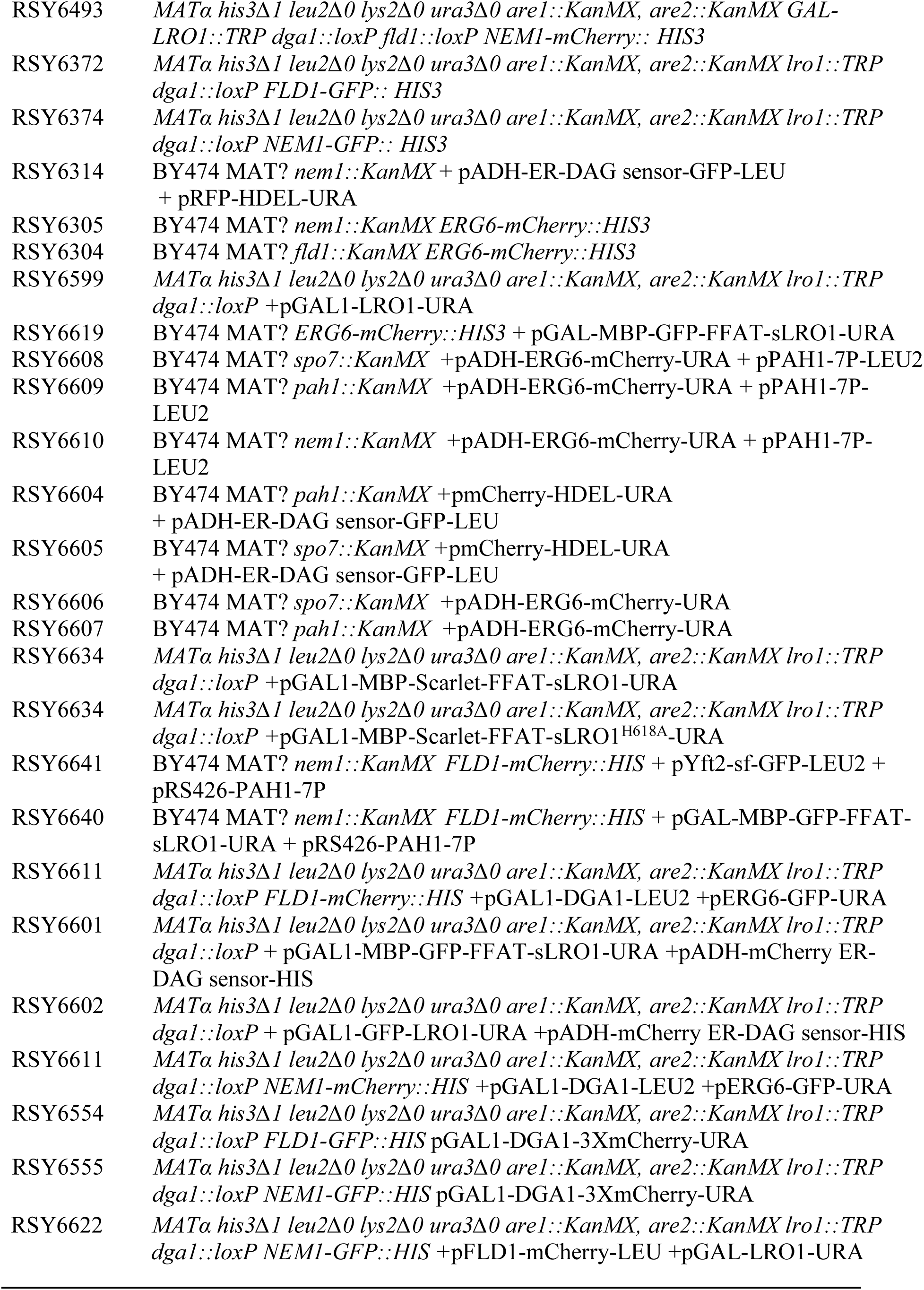
*S*.*cerevisiae* strains and plasmids used in this study.

sfGFP, superfold GFP

Yft2-sfGFP is expressed under GPD promoter on a CEN/LEU2 plasmid

Sec63-GFP is expressed under SEC63 promoter on a CEN/LEU2 plasmid

GFP-FFAT is expressed under GAL1 promoter on a 2µ/URA3 plasmid

GFP-LRO1 is expressed under GAL1 promoter on a 2µ/URA3 plasmid

GFP-sLRO1(98-661) is expressed under GAL1 promoter on a 2µ/URA3 plasmid ssKar2-GFP-sLRO1(98-661)-HDEL is expressed under GAL1 promoter on a 2µ/URA3 plasmid

MBP-GFP-FFAT-sLRO1(98-661) is expressed under GAL1 promoter on 2µ/URA3 plasmid

MBP-Scarlet-FFAT-sLRO1(98-661) is expressed under GAL1 promoter on 2µ/URA3 plasmid

MBP-Scarlet-FFAT-sLRO1^H618A^ is expressed under GAL1 promoter on 2µ/URA3 plasmid

FLD1-mCherry is expressed under ADH1 promoter on a CEN/LEU2 plasmid

ERG6-GFP is expressed under native promoter on a CEN/URA3 plasmid

mCherry-HDEL is expressed under ADH1 promoter on a CEN/LEU2 plasmid

RFP-HDEL is expressed under ADH1 promoter on a CEN/LEU2 plasmid

ER DAG sensor (PKD-GFP-Ubc6) is expressed under ADH promoter on a 2µ/LEU2 plasmid

## Notes

### Competing Interest Statement

The authors have declared no competing interest.

